# Intracellular growth of *Chlamydia trachomatis* leads to global histone hypermethylation by impairing demethylation

**DOI:** 10.1101/2024.06.04.597420

**Authors:** Chloé I. Charendoff, Félix V. Louchez, Yongzheng Wu, Lee Dolat, Guillaume Velasco, Stéphanie Perrinet, Adrian Gabriel Torres, Laure Blanchet, Magalie Duchateau, Quentin Giai Gianetto, Mariette Matondo, Laurence Del Maestro, Slimane Ait-Si-Ali, Frédéric Bonhomme, Gaël A. Millot, Vannary Meas-Yedid, Lluís Ribas de Pouplana, Elisabeth D. Martinez, Raphael H. Valdivia, Agathe Subtil

**Author notes:** Equal contribution (Chloé I. Charendoff, Félix V. Louchez).

## Abstract

*Chlamydia trachomatis*, an intracellular bacterium, highjacks metabolites from the host cell for its own proliferation. We provide evidence of global hypermethylation of the host proteome, including histones, during the late stages of infection. Single cell analyses revealed co-occurrence of several methylated residues on histones. Histone hypermethylation correlated with bacterial load and was prevented by antibiotic treatment. Mapping of trimethylation of histone 3 at residues K4 and K9 revealed a broad distribution throughout the chromatin. Nuclear fractions of infected cells exhibited a four-fold decrease of demethylase activity towards H3K4me3 and a two-fold increase in succinate concentration, a competitive inhibitor for the demethylase co-factor α-ketoglutarate. Supplementation of the culture medium with dimethyl-ketoglutarate (DMKG) or with iron, a second co-factor of histone lysine demethylases, reduced histone hypermethylation. DMKG supplementation modified the transcription of about one third of the infection-responsive genes, indicating that histone hypermethylation contributes to modulating the transcriptional response of the host to infection. Finally, chemical inhibition of histone demethylases in a mouse model of infection showed a moderate benefit regarding the outcome of infection. Overall, our data show that the metabolic pressure exerted by a pathogen with an intracellular lifestyle drives epigenetic changes in infected cells.

## INTRODUCTION

Chemical modifications of histones or DNA determine chromatin condensation, DNA accessibility and recruitment of the transcription machinery. Methylation of DNA and histones are the best characterized chromatin modifications. In general, DNA methylation is associated with chromatin compaction and inhibition of transcription, while histone methylation can either promote or repress transcription depending on the specific histone and lysine residues modified (Luger et al., 2012).

Chromatin dynamics is regulated by cellular metabolism as several metabolites support chromatin remodeling. All methylation reactions rely on the availability of S-adenosyl methionine (SAM) (Su et al., 2016) and numerous chromatin modifiers use metabolites as cofactors (Boon et al., 2020). The enzymatic activity of many histone lysine demethylases requires the 2-oxoglutarate oxygenase activity of a Jumonji C (JmjC) domain with 2-oxoglutarate (also known as α-ketoglutarate; aKG) and iron II (Fe(II)) needed for the demethylation reaction (Accari and Fisher, 2015). The activities of the histone demethylases are sensitive to physiological variations in the concentration of these metabolites (or of competitive metabolites), and thereby metabolic changes can influence epigenetic modifications (Jiang et al., 2019; Shapiro et al., 2023; Xiao et al., 2012). Overall, the effect of metabolites on the epigenetic status of chromatin are well documented in metabolic diseases and in cancer (Boon *et al*., 2020; Reid et al., 2017). Metabolites are also at the heart of the influence of the mammalian gut microbiota on host developmental, immunologic, and metabolic outcomes (Abu Kwaik, 2015; Krautkramer et al., 2016; Nichols and Davenport, 2021; Zhao et al., 2021).

The obligate intracellular bacterium *Chlamydia trachomatis* infects epithelial cells of the genital tract and of the eye conjunctiva (Brunham and Rey-Ladino, 2005). The bacteria undergo a biphasic developmental cycle, which takes place entirely within a vacuolar compartment, called an inclusion, and lasts about 48 h *in vitro* (AbdelRahman and Belland, 2005). All metabolites required for bacterial growth are captured from the host cytoplasm. A decade ago, a seminal study revealed that several histone methylation marks were increased in epithelial cells infected with *C. trachomatis* (Chumduri et al., 2013) but the mechanism underlying this shift in chromatin dynamics during infection is unknown. Furthermore, histone modifications normally associated with opposite outcomes in terms of gene expression, such as H3K4me3 and H3K9me3, were similarly upregulated at the population level, so that the global outcome of these changes on host transcription was not predictable.

Here, we characterized the nature of histone modification during *C. trachomatis* infection at the single cell level. We observed that the level of histone methylation was variable between infected cells, and that the proportion of cells that displayed a high level of histone methylation increased with infection time. Strikingly, at the individual cell level, the levels of methylation of different lysine residues strongly correlated with each other. At the proteome level we saw an overall increase in protein methylation in infected cells. We then undertook to understand the origin of histone hypermethylation in *Chlamydia* infected cells, the genomic distribution of two of these epigenetic modifications and whether histone hypermethylation contributed to the transcriptional response of the host to infection.

## RESULTS

### *C. trachomatis* infection induces protein hypermethylation including on histones

Several histone methylation modifications (H3K4me3, H3K9me3, H3K27me3, H3K79me2, H4K20me3) were examined by immunofluorescence in cells infected for 48 h with *C. trachomatis* serovar LGV L2 constitutively expressing the fluorescent protein mCherry. All modifications displayed a strong signal in several, but not all, infected cells (Fig. 1A and S1A). Strikingly, most cells positive for one modification were also positive for a second modification. An automated procedure to quantify fluorescence intensities in nuclei confirmed the increase in methylated histone lysine residues in nuclei of infected cells compared to non-infected ones of the same coverslip. When co-stained residues were examined a positive correlation between the intensities was observed (Fig. 1A). Histone lysine hypermethylation was not restricted to *C. trachomatis*, as the mouse adapted strain *C. muridarum* also induced histone hypermethylation late in infection in HeLa cells and in mouse embryonic fibroblasts (MEFs) (Fig. S1B-C). Histone hypermethylation in MEFs was also observed upon infection with *C. trachomatis* (Fig. S1C).

**Fig. 1.**
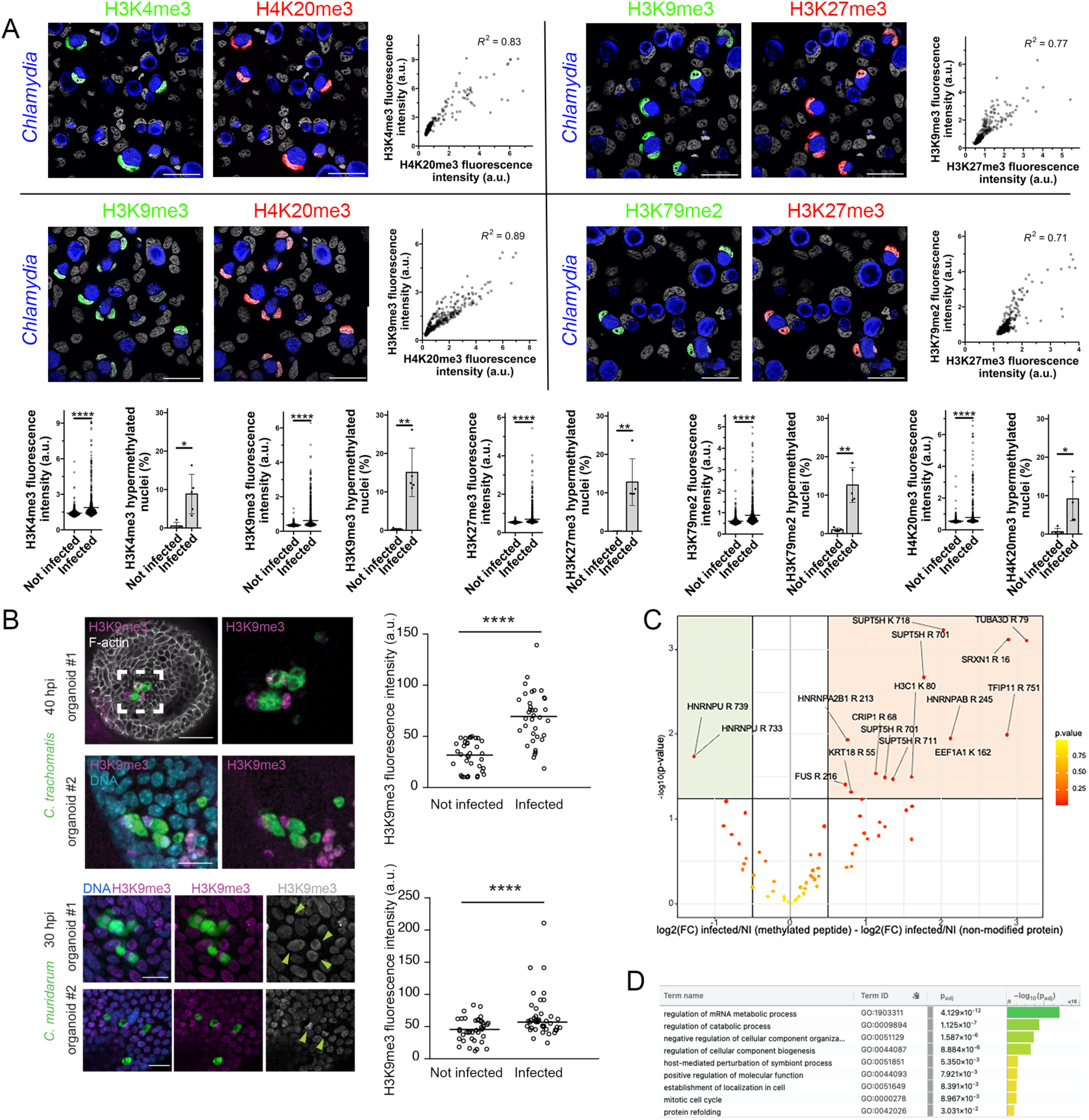
*C. trachomatis* infection induces protein hypermethylation. **A**. Cells were infected at MOI=1 with *C. trachomatis* serovar L2 for 48 h before fixation, permeabilization and staining with the indicated antibodies. DAPI staining of DNA allows for the detection of host nuclei and bacteria. *Chlamydia* inclusions are circled for reference. Bar, 50 μm. Mean intensities for the indicated methylated residues were quantified in the nuclei of non-infected and infected cells of the same coverslip. a.u., arbitrary unit. Histograms display the percentage of positive nuclei for the indicated histone methylation mark, defined as nuclei with fluorescence intensity greater than 2-fold the mean intensity of nuclei of non-infected samples. The p-value of a Welch’s t-test is shown ****: P < 0.0001, **: P < 0.01, *: P < 0.05 R= Pearson’s coefficient of correlation. **B**. Mouse derived organoids infected for 40 h with GFP-expressing *C. trachomatis* serovar L2 (top) or for 30 h with GFP-expressing *C. muridarum* (bottom) were fixed and stained for H3K9me3 (purple). Arrow heads point to nuclei with increased H3K9me intensity. Quantification of the intensity of the signal in nuclei of infected cells and of neighboring non infected cells is shown. The p-value of a Welch’s t-test is shown. Bars, 20 μm. **C**. Volcano-plot of the relative enrichment of methylated host proteins in cells infected with *C. trachomatis*. The site of methylation (on R or K) is indicated for the proteins that are significantly hypomethylated (green quadrant) or hypermethylated (orange quadrant) in infected cells. Proteins or methylated peptides that were recovered in only one condition are absent from this representation (Table S1). D. Analysis of the biological pathways associated to proteins differentially methylated late in infection using g:Profiler (Kolberg et al., 2023).

To validate these observations in non-cancer derived cells we first used primary epithelial cells derived from the ectocervix of female patients (Tang et al., 2021). The number of nuclei positive for the three histone methylation modifications tested (H3K9me3, H3K27me3, H4K20me3) was higher in the infected cells compared to non-infected cells (Fig. S1D). Finally, we used a mouse endometrial organoid model that recapitulates epithelial cell diversity, polarity and ensuing responses to *Chlamydia* infection (Dolat and Valdivia, 2021). We observed an increase in H3K9me3 intensity in nuclei of infected cells, compared to the surrounding non-infected cells, when organoids were infected with *C. trachomatis* or *C. muridarum* (Fig. 1B). These observations confirm the ubiquity of global histone hypermethylation in the late stages of infection.

To understand whether histone post-translational modifications (PTMs) late in infection was specific to histones or the result of a general increase in protein methylation, we applied mass spectrometry-based bottom-up proteomics to assess the level of methylation of proteins in infected versus uninfected cells. Statistical analyses were performed on confidently identified methylated peptides encompassing mono-, di-, or tri-methylations. To control for variations in the relative abundance of proteins because of infection, the changes in quantified methylation values were normalized to the abundance of their cognate proteins, quantified from non-modified peptides. These analyses revealed an increase in protein methylation in infection on arginine and lysine residues (Fig. 1C and Table S1). Despite the inherent challenges associated with the combinatorial nature of post-translational modifications on histones, we detected hypermethylation of H3K79 in infected samples. In addition, we observed increased methylation in several proteins including proteins implicated in mRNA synthesis (e.g. SUPT5H, TFIP11, HNRNPAB, HNRNPA2B1) and the cytoskeleton (TUBA3D and KRT18). Overall, gene ontology analysis indicated that mRNA metabolism was the biological pathway that counted the highest proportion of proteins differentially methylated late in infection (Fig. 1D). In addition, although the abundance of some of the known methyl transferases and demethylases recovered in the proteomic data varied in abundance between infected and not infected samples there was no global trend that could account for the global increase in methylation (Table S2).

Finally, to assess whether hypermethylation extended to host DNA, we quantified the methylation level of cytosine residues in nuclei from infected vs. non-infected cells by LC-MS/MS analysis. In contrast to what we had observed on histones, host DNA was slightly hypomethylated in infected cells (Fig. S1E). Altogether, we concluded that the hypermethylation of histones during *C. trachomatis* infection occurred in the context of global protein, not DNA, hypermethylation.

### Histone hypermethylation is linked to bacterial metabolic activity but not to the bacterial effector Nue

We next analyzed the kinetics of occurrence of histone hypermethylation using two representative marks, e.g. H3K9me3 and H4K20me3. Histone hypermethylation was rarely observed in the first twenty-four hours of the infectious cycle, indicating that it occurs in the second half of the infectious cycle, when bacterial load is high (Fig. 2A). To further link the level of histone modification with bacterial burden, we used the level of fluorescence of mCherry as a readout for bacterial load. We observed that, within the infected cells, the sum intensity of the mCherry signal was higher in cells that displayed hypermethylation of H3K9me3 than in cells with low level of H3K9me3, indicating that histone hypermethylation correlated with bacterial load (Fig. 2B). If true, slowing down bacterial growth with the addition of an antibiotic at mid-course should reduce the methylation phenomenon late in infection. We chose to use the fluoroquinolone ciprofloxacin as it does not affect mitochondrial activities in the host (Riesbeck et al., 1990). Addition ciprofloxacin in the last 24 h of infection reduced bacterial load and the level of histone lysine methylation (Fig. 2C). Altogether, we concluded that histone hypermethylation required sustained bacterial proliferation.

**Fig. 2.**
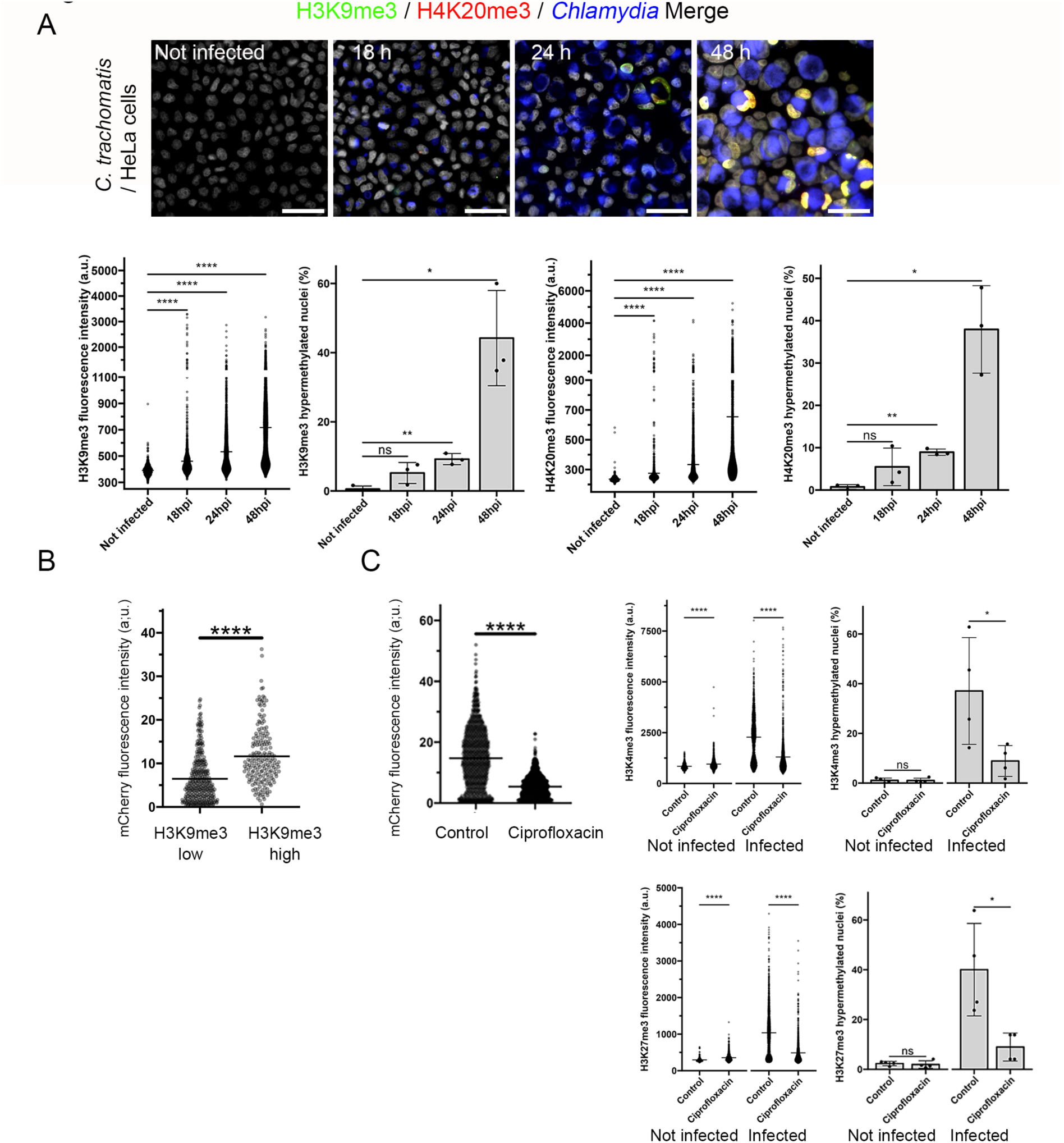
Histone hypermethylation correlates with bacterial proliferation. **A**. HeLa cells were infected with mCherry expressing *C. trachomatis* serovar L2 (MOI=1) for the indicated time before fixation, permeabilization and staining for H3K9me3 and H4K20me3. The mCherry signal is displayed in blue, DNA in white. Bar, 50 μm. Dot plots display the mean intensities for the indicated methylated residues in one representative experiment. a.u., arbitrary unit. Histograms display the percentage of positive nuclei for the indicated histone methylation mark, defined as nuclei with fluorescence intensity greater than 2-fold the mean intensity of nuclei of non-infected cells. Data from three independent experiments and the p-values of Welch’s t-tests are shown ****: P < 0.0001, **: P < 0.01, *: P < 0.05. **B**. HeLa cells were infected with *C. trachomatis* L2 expressing mCherry for 48 h before fixation, permeabilization and staining for H3K9me3. The mean fluorescence intensity of mCherry of each individual inclusion is shown for infected cells with a low level of H3K9me3 signal (signal < 2-fold mean H3K9me3 signal in nuclei of non-infected cells) and with a high level of H3K9me3 signal (signal > 2-fold mean H3K9me3 signal in nuclei of non-infected cells). The p-value of a Welch’s t-test is shown. **C**. Cells were infected with mCherry expressing *C. trachomatis* serovar L2 and 15 μg/ml ciprofloxacin was added to the culture medium 24 hpi. Cells were fixed 48 hpi, permeabilized and stained for H3K4me3 and H3K27me3. The dot plots display the mean mCherry intensities in inclusions of control and ciprofloxacin treated cultures (*left*) and the relative mean intensities of H3K4me3 (*top*) and H3K27me3 (*bottom*) in the nuclei from infected cells and non-infected cells in one representative experiment. Histograms display the percentage of positive nuclei in four independent experiments. The p-values of Welch’s t-tests are shown.

We next asked whether Nue, a protein secreted by *Chlamydia* endowed with histone methyl transferase activity (Pennini et al., 2010), was implicated in this process. We obtained a *C. trachomatis* strain with a nonsense mutation in *nue* (C to T mutation at position 129786 leading to a W71* truncation), designated as CTL2M340 (Kokes et al., 2015) and confirmed the absence of Nue expression by western blot analysis (Fig. S1F). Histone hypermethylation was still observed in cells infected for 48 h with CTL2M340, demonstrating that Nue is not required for histone hypermethylation observed late in infection (Fig. S1F).

### Histone hypermethylation is the result of reduced demethylase activity late in infection

The histone methylation modifications we observed in infected cells are deposited by several types of histone methyl transferases and removed by an equally diverse set of dedicated demethylases (Accari and Fisher, 2015; Husmann and Gozani, 2019; Maiques-Diaz and Somervaille, 2016). Because some of these enzymes use common co-factors we reasoned that changes in the concentrations of these co-factors due to infection could affect several histone modifications simultaneously. Hypermethylation could be the result of increased methylation or decreased demethylase activity. Since all methylation reactions use SAM as a co-factor, we determined SAM concentrations from isolated host nuclei using a fluorescence transfer-based assay. We observed a 50 % decrease in the amount of nuclear SAM in cells infected by *C. trachomatis* for 48 h, suggesting that histone hypermethylation is not a consequence of SAM accumulation during infection (Fig. 3A). We thus considered the alternative hypothesis that histone demethylase activity may decrease late in infection. To test it, we first looked at whether a demethylase inhibitor could reproduce the histone hypermethylation phenotype. JIB-04 is a membrane permeable small inhibitor of all JmjC demethylases (Wang et al., 2013). When the inhibitor was added to the culture medium 24 h after infection by *C. trachomatis*, and the cells fixed and stained 48 hpi, we observed a dose-dependent increase in the H4K20me3 positive cells among the infected cells (Fig. 3B). A similar result was obtained for H3K9me3 (Fig. S2A-B). The increase in histone methylation induced by JIB-04 treatment was much more pronounced in the infected than in the non-infected cells. Note that hypermethylation occurred despite a negative effect of JIB-04 treatment on bacterial development (Fig. S2C). Other demethylase inhibitors (SD70 and TACH101) also increased the methylation of their target histone lysine H3K9 (Fig. S2D), while impairing bacterial replication (Fig. S2E). These data indicate that infected cells are sensitized to the action of inhibitors of JmjC demethylases, consistent with the hypothesis that lysine demethylation by JmjC demethylases is low late in infection. If this hypothesis is correct, overexpression of a given JmjC demethylase should reverse the enhanced methylation of its target residue. KDM4A (also known as JMJD2A) is a demethylase specific for H3K9me3 and H3K36me3, while KDM5A (JARID1A) demethylates H3K4me3 to di- and mono-methylated forms (Iwase et al., 2007; Whetstine et al., 2006). Cells were infected with mCherry expressing *C. trachomatis* L2 and transfected 2 hours later with HA-tagged demethylases, or catalytically dead enzymes as negative controls. We observed a 60% reduction in the proportion of infected cells that displayed H3K4me3 hypermethylation in cells expressing HA-KDM5A, but not the catalytically dead version of the enzyme. In contrast, the proportion of infected cells positive for H3K9me3 was unchanged upon overexpression of KDM4A. Overexpression of KDM4A had the opposite effect: it reduced the proportion of cells displaying H3K9me3 hypermethylation without affecting H3K4me3 levels. Overexpression of the catalytically dead KDM4A mutant affected neither H3K4me3 nor H3K9me3 levels (Fig. 3C). Thus, overexpression of a demethylases was sufficient to partially prevent hypermethylation of its specific target. This result is fully consistent with the hypothesis that hypermethylation results from low activity of histone demethylases, that can be specifically overcome by overexpression of a cognate demethylase. Finally, we directly measured the demethylase activity in nuclear extracts of infected and non-infected cells against one specific substrate, i.e. trimethylated H3K4. The demethylase activity in nuclear extracts from infected cells was reduced to about 25% of its level in nuclear extracts of non-infected cells, supporting the hypothesis of a global decrease in histone demethylase activity in the host nucleus late in infection (Fig. 3D).

**Fig. 3.**
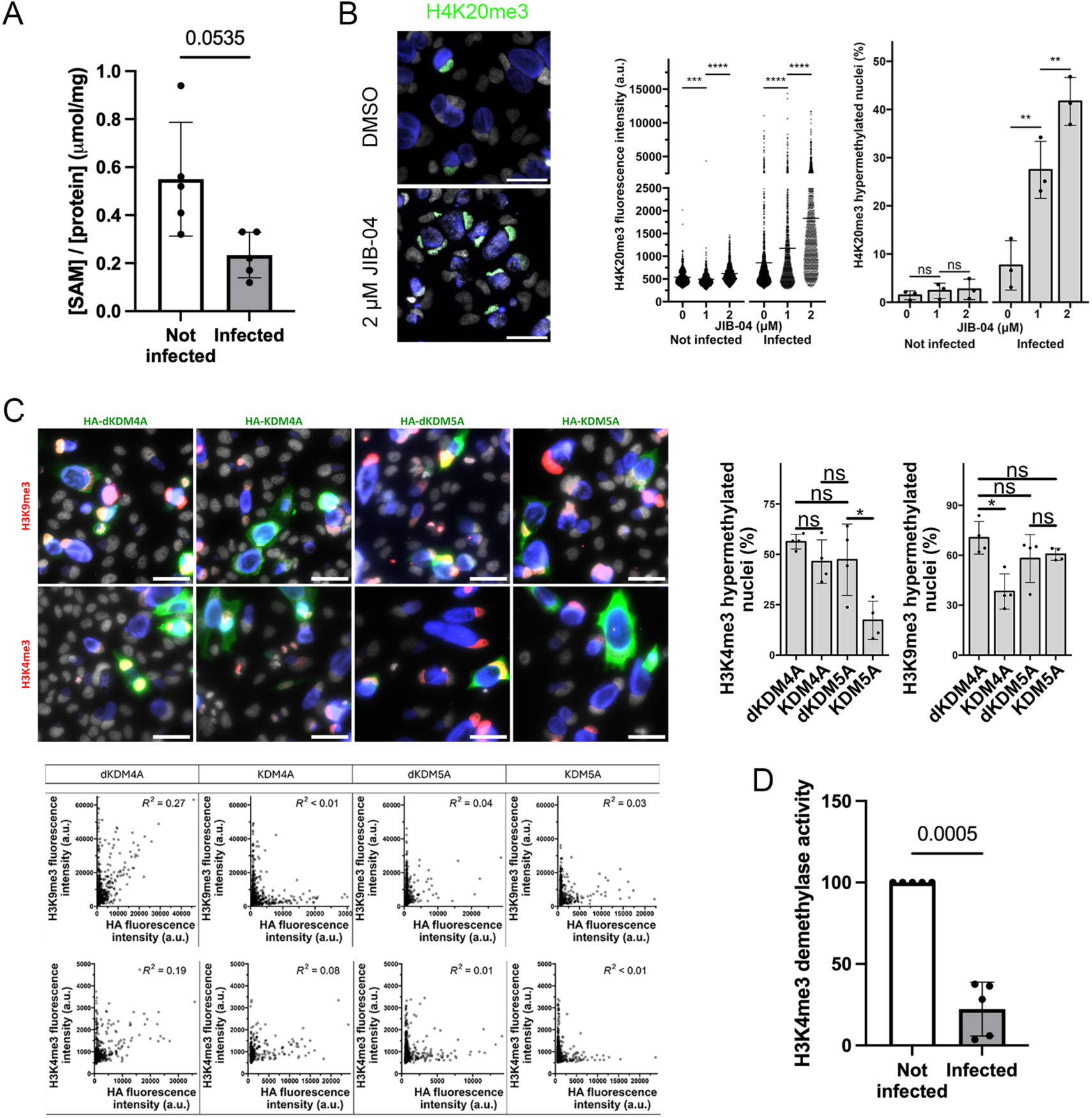
The activities of histone demethylases are low in late *C. trachomatis* infected cells. **A**. Cells were infected with *C. trachomatis* serovar L2 for 48 h before isolation of the nuclei. SAM concentrations were normalized to protein content. Each dot represents an independent experiment, the mean ± SD is shown. The p-value of a ratio paired t-test is indicated. **B**. Cells were infected with mCherry expressing *C. trachomatis* serovar L2. After 24 the culture medium was refreshed with or without the indicated concentration of JIB-04. Cells were fixed 48 hpi, permeabilized and stained for H4K20me3 (green). DNA was labeled with DAPI (white), mCherry is displayed in blue. Bar, 50 μm. The dot plot displays H4K20me3 mean fluorescence intensity in nuclei of non-infected or infected cells. a.u., arbitrary unit. Black bars indicate the mean. The p-value of Welch’s t-test are shown. The histogram displays percentage of positive nuclei in three independent experiments. The p-value of Welch’s t-tests are shown ****, P < 0.0001; ***, P < 0.001; **, P < 0.01; *, P < 0.05; n.s., not significant. Nuclei were scored positive when H4K20me3 intensity was greater than two times its mean intensity in nuclei of non-infected cells from the same culture condition. **C**. Cells were transfected with the indicated HA-tagged constructs (dKDM: catalytical dead mutant) and infected 3 h later with mCherry expressing *C. trachomatis* L2. Cells were fixed 48 h later, permeabilized and stained for HA and for the indicated methylated residues. DNA was labeled with DAPI (white), mCherry is displayed in blue. Bar, 50 μm. The dot plots display mean fluorescence intensities of the indicated marks in one representative experiment, each dot representing one infected cell. The histograms display the percentages of positive nuclei in the transfected infected cells in four independent experiments. Cells were considered transfected when the HA signal in the nucleus was greater than the two-fold value of the mean intensity in nuclei in non-transfected cells of duplicate wells. Nuclei were scored positive as in B. **D**. Demethylase activity against H3K4me3 was measured in nuclear extracts from cells infected for 40 h, normalized to protein concentration and expressed relative to the activity measured in nuclear extracts from non-infected cells. The p-value of a Welch’s t-test is indicated.

### Supply of demethylase cofactors reduces histone hypermethylation

JmjC demethylases require oxygen, iron (FeII) and aKG, and a limitation of one or more of these three co-factors could impair their activity. Increase in succinate and fumarate levels could also account for the defect in histone demethylase activity, as those are competitive inhibitors of the 2-oxoglutarate oxygenase activity carried out by Jumonji C domains (Xiao *et al*., 2012). Oxygen can be excluded, as the experiments were performed under normoxic conditions. Interpretation of the measurements of metabolite concentrations in infected cells is obscured by the contribution of metabolites located in the bacteria, which, late in infection, represent a significant fraction of the total metabolites. To circumvent this difficulty and focus on the compartment in which the demethylase activity occurs, we measured succinate concentrations in nuclear fractions, devoid of bacteria. We observed a two-fold increase in the concentration of succinate in nuclei isolated 40 hpi compared to control nuclei (Fig. 4A). This result supports the hypothesis that a decrease in the aKG to succinate ratio during infection contributes to the loss in histone demethylase activity. If iron and/or aKG became limited during infection, supplementation of the culture medium with these metabolites might restrain this phenomenon, provided they are transported to the host cytoplasm. Iron was supplemented with addition of 100 μM ferric ammonium citrate in the culture medium, a concentration sufficient to reverse iron deficiency *in vitro* (Jiang *et al*., 2019). For aKG we supplemented the culture medium with 5 mM dimethyl-ketoglutarate (DMKG), which is membrane permeable and converted into aKG by cytosolic esterases. Interestingly, we observed that changing the medium at mid-course during infection (24 hpi) was sufficient to partially reduce H3K4 and H4K20 tri-methylation at 48 hpi, possibly by replacing rate-limiting metabolites depleted during infection (Fig. 4B-C). Supplementation of iron or DMKG reduced the level of histone methylation even further. To control for the possible effect of these treatments on bacterial growth, we measured their impact on the infectious forming units (IFUs) produced within 48 h. Iron supplementation had no effect on IFUs while DMKG supplementation reduced the infectious progeny by about 25% and combining iron and DMKG reduced progeny by about 50% (Fig. S2F). The effect of the combination was not tested as the reduction on bacterial growth could be sufficient on its own to prevent histone methylation. Altogether, these experiments show that DMKG or iron supplementation of the culture medium diminishes hypermethylation of histone lysine residues late in infection. Together with the observation that succinate levels increase in infection, the data support the hypothesis that the reduction in JmjC histone demethylase activities is driven by unbalance in the concentration of their co-factors.

**Fig. 4.**
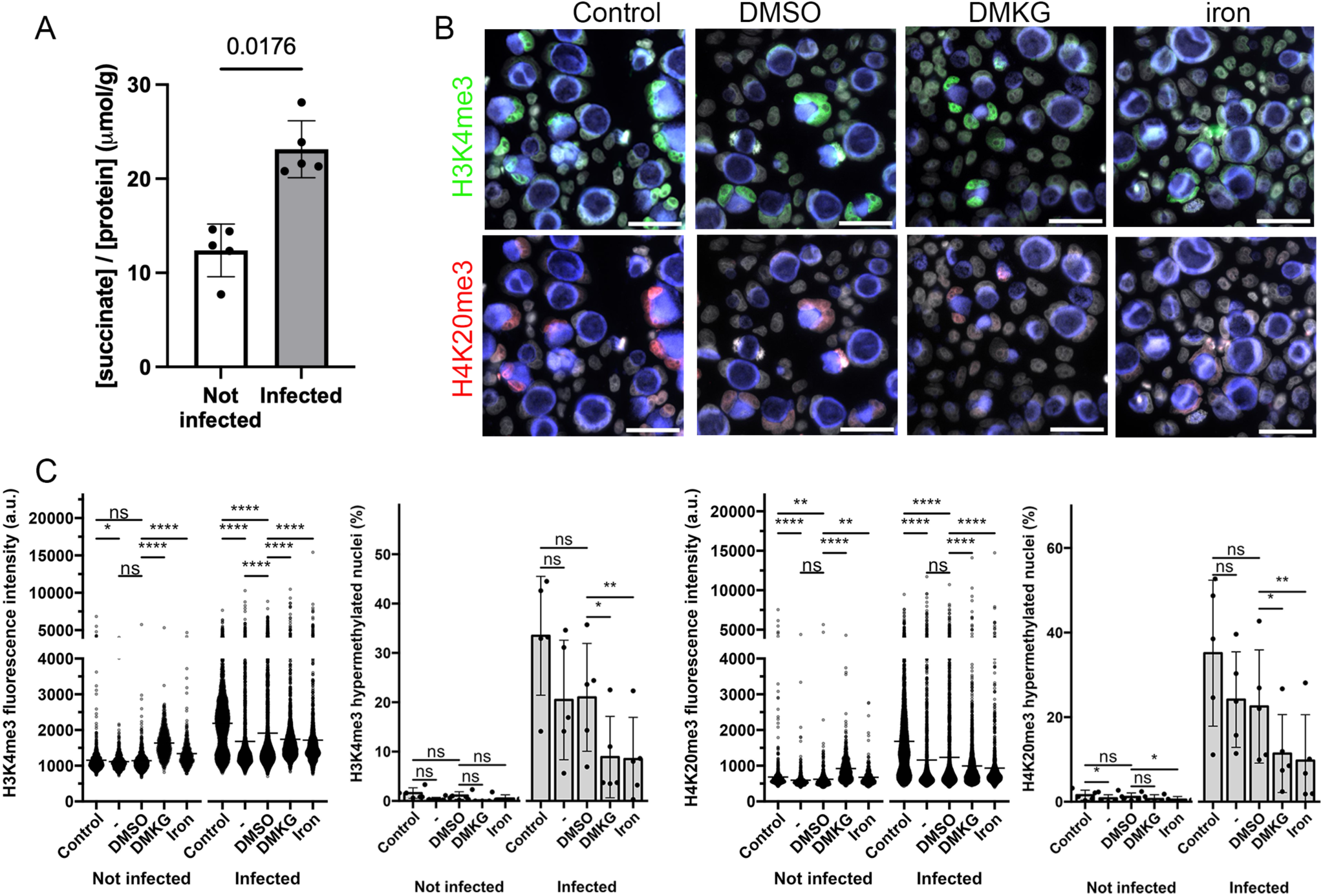
Supply of demethylase cofactors reduces histone hypermethylation. **A**. Succinate concentrations were measured in nuclear extracts from cells infected for 40 h or not infected and normalized to protein concentrations. The p-value of a ratio paired t-test is indicated. **B**. Cells were infected with mCherry expressing *C. trachomatis* serovar L2 (MOI=2). After 24 h the culture medium was left untouched (control) or refreshed with the addition of 5 mM DMKG, solvent alone (DMSO) or 100 μM ferric ammonium citrate (iron). Cells were fixed 48 hpi, permeabilized and stained for H3K4me3 (green) and H4K20me3 (red). DNA was stained with DAPI (display in white). Bar, 50 μm. **C**. The dot plots show the intensities for the indicated methylated residues in nuclei of infected and non-infected cells for one representative experiment. The second condition corresponds to refreshment of the medium without addition of DMSO. The p-value of Welch’s t-tests are shown. The histograms display the percentages of positive nuclei in 5 independent experiments. The p-value of Welch’s t-tests are shown.

### Hypermethylation on H3K4 and H3K9 spreads throughout chromatin

Does histone hypermethylation occur randomly across chromatin, or are some genomic regions more exposed? To answer this question, we investigated the pattern of chromosomal deposition of two well-studied histone methylation modifications, H3K4me3 and H3K9me3, associated with active and repressed chromatin, respectively, in infected cells by chromatin immunoprecipitation followed by DNA sequencing (ChIP-seq). We observed an increase in the amount of material immunoprecipitated from infected samples compared to non-infected samples, consistent with the increase of these methylation modifications during infection (Fig. S3A). Samples were adjusted to equal DNA concentration before sequencing. Peak calling showed that the number of peaks for H3K4me3 or H3K9me3 were not markedly changed (Fig. S3B). H3K4me3 was enriched in active promoters whereas H3K9me3 was enriched in heterochromatin and regions of low activity, as expected. The patterns of the distribution of the marks across the different genomic regions were globally unchanged between infected and non-infected samples, suggesting that hypermethylation did not accumulate preferentially in specific chromatin regions for these two marks (Fig. 5A). Since the two independent experiments showed the expected hierarchical clustering (Fig. S3C) we merged the reads from biological replicates pairs for H3K4me3 and H3K9me3 for further analysis. The density of reads in H3K4me3 peaks diminished in infected samples (Fig 5B, Table S3) and was accompanied by a slight but significant increase of the median of read coverage all over the genome in the immunoprecipitated fractions and not in the input (Fig. 5C). These data are consistent with H3K4 hypermethylation in infected cells occurring outside H3K4me3-enriched chromatin rather than reinforcing existing peaks. In other words, the inhibition of H3K4me3 demethylation increases the proportion of methylated H3K4 outside H3K4me3-enriched areas of chromatin. The median of read coverage all over the genome also increased slightly in the H3K9me3 immunoprecipitated fractions (Fig. 5C). Only 20 sites were significantly differentially enriched for H3K9me3 between the two conditions (Table S3). These data suggest that, while globally increasing late in infection, the pattern of deposition of H3K9me3 remained similar to that of non-infected cells. Altogether, histone hypermethylation did not appear preferentially in specific chromatin regions, at least for the two marks investigated. The data were consistent with a random reduction of demethylase activity across host chromatin.

**Fig. 5.**
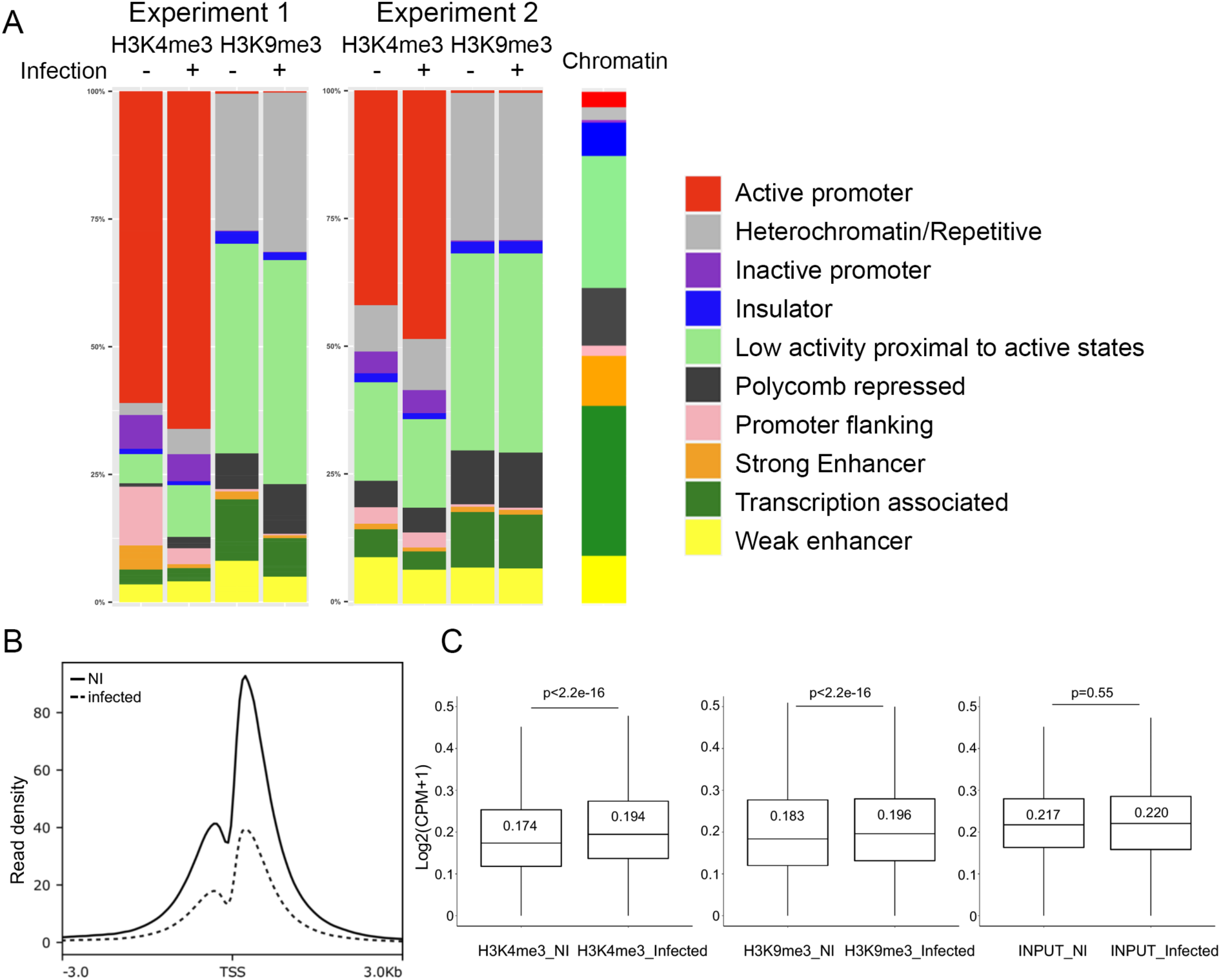
H3K4me3 and H3K9me3 chromatin landscape. **A.** Distribution of genomic sites enriched in H3K4me3 and H3K9me3 marks. The functional annotation was based on the chromatin states established from ChromHMM (ChromHMM track from ENCODE/Broad Institute). The relative abundance in HeLa cells of each chromatin state is shown on the right. **B**. Profile of the H3K4me3 ChIP-seq signal (read density) from merged biological replicates, i.e pileups of the H3K4me3 enrichment relative to chromatin input (subtraction mode, input from IP of bam files) in non-infected (NI) and infected HeLa cells on genomic loci defined as 3kb upstream and downstream of annotated Transcriptional Start Site (TSS) of genes (NCBI RefSeq). **C**. Boxplots showing reads coverage (log2(CPM+1); CPM, counts per million; bin=500 bp) from H3K4me3, H3K9me3 ChIP-seq experiments and input in non-infected (NI) and infected HeLa cells. The median and the p-value (Wicoxon test) is indicated within each boxplot. Outliers are not shown for clarity.

### DMKG supplementation modifies the host cell’s transcriptional response to infection

Histone methylation can either promote or repress transcription depending on the specific histone and lysine residues modified (Luger *et al*., 2012). Given that *Chlamydia* infection resulted in epigenetic modifications associated with antagonist outcomes we wondered whether histone hypermethylation globally shifted the transcriptional host response to infection in a given direction. To answer this question, cells were infected with or without supplementing the culture medium with DMKG at 24 hpi to restrain histone hypermethylation. Non-infected cells were treated in parallel. RNA was extracted from these cells 48 hpi and processed for analysis by microarray. The differentially expressed genes (DEGs) were defined with a cut-off of |Fold-increase|>2 and a p-value <0.05. The number of elements differentially up or down regulated between any conditions are split evenly (Fig. 6A). In non-treated samples, we observed 1055 genes were differentially regulated by infection (Fig. 6A and Table S4). About one third of those (364) were no longer differentially expressed when DMKG was added to the culture medium, indicating that increasing intracellular levels of aKG during infection impacts the host cell’s transcriptional response to infection (Fig. 6B).

**Fig. 6.**
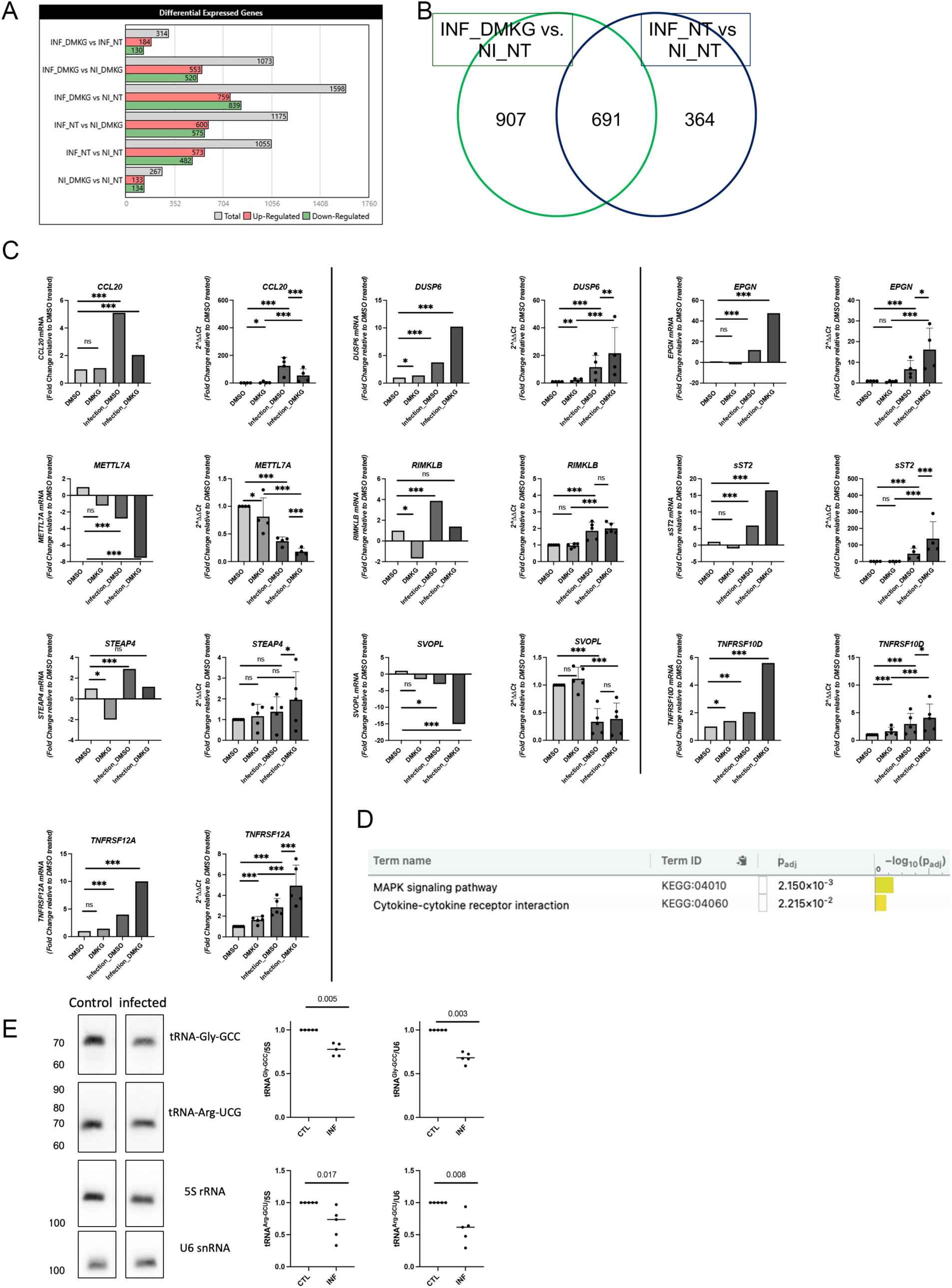
DMKG supplementation modifies the host transcriptional response to infection. **A**. Transcripts from infected (INF) and non-infected (NI) samples, supplemented or not with 5 mM DMKG 24 hpi, and extracted 48 hpi were analyzed by microarray. Two by two comparisons of differentially expressed genes are shown (cut-off: 2-fold difference). **B**. Venn diagram showing about one third (364/1055) of the differentially regulated genes upon infection were no longer differentially regulated with DMKG supplementation. **C**. For each of the ten genes tested, the histogram on the left displays the microarray result (p_values were extracted from Table S5) and the histogram on the right shows the result of the RT-qPCR validation. For RT-qPCR validation samples were prepared in the same conditions as the microarray samples. Specific sets of primers for the indicated genes were used to measure transcripts levels by RT-qPCR normalized to *actin* transcripts following the ΔΔCt method. Each experiment was performed repeated at least four times (dots). The data are presented as relative mRNA levels compared to uninfected cells and shown as the mean ± SD. Statistical comparisons were obtained using contrast analysis after linear modeling, 2 by 2 multiple-testing and P-value adjustment (See methods for details). ****, P < 0.0001; ***, P < 0.001; **, P < 0.01. **D**. KEGG pathways associated to the 55 infection responsive genes differentially regulated by DMKG supplementation (Table S5 tab 3, analysis with g:Profiler). **E**. Cells were infected with *C. trachomatis* L2 for 48 hpi. Levels of tRNA^Gly-GCC^ and tRNA^Arg-UCG^ measured by northern blot are expressed relative to 5S RNA and U6 snRNA. See the method section for statistical analysis of the data.

To validate these microarray data, we tested by reverse transcription quantitative PCR (RT-qPCR) the mRNA levels of the 10 coding genes that displayed the strongest response to infection (>2-fold difference between infected and non-infected samples) and to DMKG addition (>2.5-fold difference between DMKG-treated and not-treated in infected samples) (Table S5). RNA was extracted at 48 hpi from infected vs non infected cells with and without DMKG supplementation at 24 hpi and compared to uninfected controls via RT-qPCR. The RT-qPCR results recapitulated the microarray data with the exception of *STEAP4*, that was not significantly induced by infection, and *SVOPL* and *RIMKLB*, that did not respond significantly to DMKG supplementation (Fig. 6C). Thus, the microarray data were validated at 90% for the response to infection (9 genes out of 10) and 78% for the DMKG sensitivity (7 genes out of 9).

Given that the histone methylation can boost of repress transcription, depending on the methylated residue, we next analyzed whether histone hypermethylation during infection globally tilted the balance of transcription regulation in one direction or the other. In non-infected cells, DMKG supplementation resulted in differential expression of 267 genes, evenly distributed between transcripts that were upregulated (133) and downregulated (134) (Fig. 6A). In infected cells, a similar range of changes was observed, with 314 DEGs, 184 being up and 130 being down (Fig. 6A, Table S5). Among the 184 DEGs that were upregulated with DMKG supplementation in infection, 32 corresponded to transcripts that were upregulated by infection, and 62 to transcripts that were downregulated by infection (the remaining 90 elements showing less than a 2-fold difference in transcription upon infection). Therefore, the shift in transcription when comparing infection versus infection with DMKG supplementation does not have a global trend. In other words, DMKG supplementation enhances the transcription of some of the genes that were already upregulated by infection (e.g. *DUSP6*), while it dampens the transcription of others (e.g. *CCL20*). The same observations were made on genes downregulated by infection. Thus, overall, the effect of DMKG supplementation on the expression of a given transcript depends on the transcript. Histone hypermethylation does not globally enhance nor repress the host response to infection.

To minimize the intrinsic effect DMKG has on transcription, we excluded from the 314 DEG list (comparison of infected and infected with DMKG supplementation, Fig. 6A) those that showed over 2-fold differential expression upon DMKG treatment alone, reducing the list to 246 DEGs. Among those, 55 correspond to genes that are differentially regulated by infection (Table S5). Analysis of this set of genes with g:Profiler pointed to a weak enrichment in genes involved in the MAPK signaling pathway (*CACNB2*, *HSPA1A*, *EFNA1*, *FGF2*, *DUSP6*, *EPHA2*) and in cytokine signaling (*IL1RL1*, *CCL20*, *TNFRSF12A*, *TNFRSF10D*, *BMPR2*) (Fig. 6D).

Finally, beside coding genes, the 246 DEGs contained many non-coding RNAs like pseudogenes, lnRNAs and piRNAs. Non-coding DEGs particularly abundant in the data set were transfer tRNAs (70 DEGs). All tRNAs in this data set were decreased in infected samples compared to non-infected samples, an effect prevented by DMKG supplementation. To confirm the depletion of tRNA late in *Chlamydia* infection by an independent approach we extracted total RNAs from uninfected cells and from cells infected for 48 h. Samples were normalized to the 18S rRNA content and analyzed by northern blot using probes specific for tRNA^Gly-GCC^ and tRNA^Arg-GCU^. Probes for U6snRNA and 5S rRNA were used to normalize the signals (Fig. 6E). The quantity of tRNA^Gly-GCC^ and of tRNA^Arg-GCU^ were reduced by about 30% in infected cells compared with non-infected cells, confirming that infection leads to a decrease in tRNA levels. Thus, our data revealed a decrease in tRNA abundance late in infection, which is antagonized by DMKG supplementation.

### Administration of JIB-04 shows limited benefit on bacterial load and tissue damages in a mouse model of *Chlamydia* infection

We have observed that exposure to JIB-04 was detrimental to bacterial development *in vitro* (Fig. S2C). The same was true for two KDM4 inhibitors tested (Fig. S2E), suggesting that chemical enhancement of histone methylation had a negative impact on bacterial proliferation. To challenge the hypothesis that demethylase inhibitors might impair bacterial development we used a vaginal *Chlamydia* infection model with the mouse adapted strain, *C. muridarum*. JIB-04 is amenable to *in vivo* studies, as it showed activity in a mouse model of cancer (Wang *et al*., 2013). We performed peritoneal injection of JIB-04 at the onset of infection (Fig. 7A). Vaginal lavages were collected on day 4 post infection to measure bacterial burden. In line with our *in vitro* observations JIB-04 administration tended to lower bacterial burden at D4 of infection (Fig. 7B). The mice were sacrificed on day 35, the genital tracts excised and their morphology examined. Consistent with a decrease in the bacterial load we observed a trend towards a reduction of the pathological manifestations of infection in JIB-04 treated animals, with less cases of hydrosalpinx (Fig. 7C) and less dilatation in the uterine horns (Fig. 7D). However, the benefit from JIB-04 treatment was limited, since severe hydrosalpinx was still observed in two of the six JIB-04 treated mice.

**Fig. 7.**
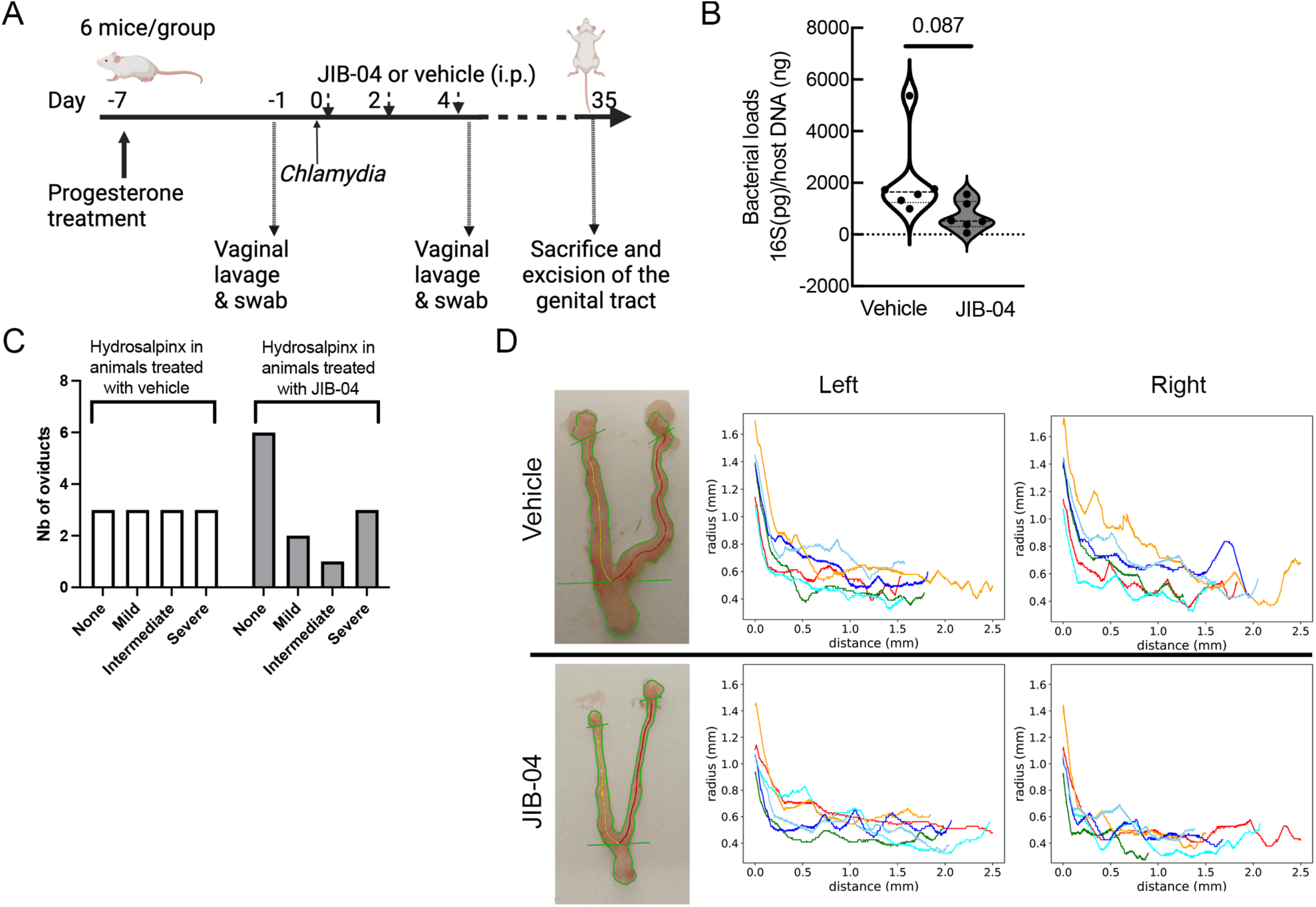
JIB-04 administration shows limited benefit *in vivo*. **A**. Schematic view of animal infections. JIB-04 treatment was administered intra-peritoneally (i.p.) and infections with *C. muridarum* were intravaginal. **B**. Bacterial load was measured on DNA extracted from vaginal lavages at D4 and normalized to host DNA. Each dot represents one mouse, the p-value of Welch’s t-test are shown. **C**. The genital tracts were excised at D35 and the presence and severity of hydrosalpinx was scored. **D** Example of the segmentation and detection of the skeletons of uterine horns. The radius along the length of the uterus horn was computed starting from the point of convergence of the right and left skeletons (yellow and red lines). Each color trace corresponds to one animal, measures of the left and right horns are displayed in left and right panels, respectively.

## DISCUSSION

Infection with an intracellular pathogen is expected to challenge the metabolic capacity of its host cell. Here we showed that infection of epithelial cells by *C. trachomatis* caused global protein hypermethylation late in infection, including hypermethylation of histone lysine residues. We provided evidence that histone hypermethylation resulted from the proliferation of bacteria and that low level of activity of host demethylases accounted for this phenomenon. Further investigation of the consequences of histone hypermethylation on the host response to infection drew a complex gene dependent picture, in line with the observation that hypermethylation led to histone modifications associated with active transcription as much as to histone modifications associated with chromatin silencing. Finally, we provide some initial evidence that drugs targeting the epigenome of the host can alter the outcome of infection *in vivo*.

Hypermethylation of several histone residues during *C. trachomatis* infection was previously documented at the cell population level (Chumduri *et al*., 2013). Our analysis by immunofluorescence showed cell to cell variability and combinatorial staining for different modifications revealed that cells that displayed a high methylation level for a given lysine also displayed hypermethylation for other residues. Hypermethylation occurred late in infection, was most prominent in cells with a high bacterial load and was prevented by antibiotic treatment, indicating that it was linked to bacterial proliferation. We excluded a role for Nue, the predicted histone methyl transferase of *C. trachomatis*, as hypermethylation was also observed in cells infected by a mutant strain that did not express Nue. A proteomic analysis of infected cells showed that, in addition to histones, several proteins were hypermethylated consistent with the observation that histone demethylases have multiple substrates (Kuwik et al., 2023). Hypermethylated proteins include several implicated in mRNA synthesis and processing, with potential consequences on host gene expression. Hypermethylation also occurred on cytoplasmic proteins such as tubulin or keratin 18. For the latter, the possibility of methylation at the site we identified had already been reported (Jang et al., 2019).

The hypermethylation of histones and its correlation with bacterial burden suggested that infection might create an environment favoring methylation reaction over demethylation. Consistent with this premise, we observed that infected cells were sensitized to the histone demethylase inhibitor JIB-04, indicating that demethylase activity is low late in infection. This was confirmed by measuring directly one such activity (on H3K4me3) in nuclei extracts isolated from infected cells. The strong loss of activity (75% reduction 40 hpi) indicated that the reduction in enzymatic activity occurred even in cells in which H3K4me3 hypermethylation was below the threshold of detection by immunofluorescence. Several observations support the hypothesis that the metabolic burden of infection is responsible for the global histone hypermethylation (*i*) histone hypermethylation increased with infection time and positively correlated with bacterial load; (*ii*) histone H3K4 and H3K9 hypermethylation was prevented by overexpression of the cognate histone demethylase (KDM5A and KDM4A, respectively); (*iii*) histone hypermethylation was partially abolished by providing aKG or Fe(II), co-factors for demethylases of the Jumonji family; and (*iv*) succinate concentration, a competitor for aKG, doubled in the nuclei of infected cells. Altogether, these data strongly suggest that aKG and Fe(II) become limiting during late infection. Indeed, Fe(II) is captured from the host cytoplasm by the bacteria (Pokorzynski et al., 2017). The host responds by enhancing expression of the transferrin receptor (3.3-fold increase of the transcript, Table S4, 5-fold increase of the protein, Table S2) but cytosolic and nuclear Fe(II) concentrations might remain at a sub-optimal concentration for demethylase activities. The second metabolite, aKG, represents an entry point into *C. trachomatis* incomplete Krebs cycle (Mehlitz et al., 2017; Stephens et al., 1998). A prior metabolomic study also detected an global increase in succinate levels in *C. trachomatis* infected cells (Rother et al., 2018), in agreement with the two-fold increase we measured in purified nuclei The same study showed that akG levels remained globally stable while fumarate, another competitive metabolite for aKG-dependent 2-oxoglutarate oxygenase (Xiao *et al*., 2012) also increased over the course of infection. Thus, increase in the ratio of succinate and fumarate over aKG, together with low iron level, can account for the decrease in demethylase activity. In agreement with the hypothesis of limiting metabolites, replenishment of the culture medium was sufficient to partially prevent histone hypermethylation. Interestingly, we show here that histone hypermethylation occurs in a context of low SAM concentration, which is required for methylation reactions. The decrease in SAM levels in infected cells was expected as *Chlamydia* highjack SAM (Binet et al., 2011). Consistent with low SAM levels, we observed that DNA was slightly hypomethylated late in infection, as previously reported (Bryan et al., 2021). How can protein hypermethylation and DNA hypomethylation be reconciled? Methylation and demethylation reactions depend on the relative affinity of each enzyme for their co-factors and on their respective rates of activities (Islam et al., 2018). This, together with the difference in turnover between DNA and histone methylation events, could account for the observed differences between DNA and histone methylation levels. In addition, enhanced expression of GADD45A and GADD45B (3.8-fold and 2.4-fold increase of the transcripts, respectively, Table S4) during infection might compensate for a potential decrease in initiation of DNA demethylation by Ten-eleven-translocation (Tet) oxidative demethylases, since GADD45 promotes DNA demethylation downstream of these enzymes (Li et al., 2015).

Epigenetic modifications govern the accessibility of chromatin. Formaldehyde-assisted isolation of regulatory elements enrichment followed by sequencing (FAIRE-Seq) experiments had shown that the proportion of all differentially accessible regions mapping to promoter regions was at its highest late in Hep2 cells infected with *C. trachomatis*, with the majority of regions showing a reduction in chromatin accessibility (Hayward et al., 2020). Our own ChIP-seq analyzes show that, at least for H3K4 and H3K9, methylation modifications are spread throughout chromatin, suggesting that hypermethylation occurs randomly. Since histone methylation can repress or activate gene transcription depending on the residue modified and its position relative to the genetic element the overall effect histone hypermethylation might have on host gene expression was unclear. To answer this question, we examined the transcriptional changes in the host response to infection upon DMKG supplementation. Interpretation of the data call for caution, since DMKG supply only partially prevents histone hypermethylation. In addition, hypermethylation affects several proteins involved in mRNA processing (Fig. 1C) and DMKG supplementation might affect transcript levels independently of its effect on histone methylation. DMKG supplementation modified the transcription of about one third of the infection-responsive transcripts, indicating that histone hypermethylation contributes to modulating the transcriptional response of the host to infection. DMKG supply did not globally enhance nor dampen the host response to infection, since similar proportion of transcripts were differentially up- and downregulated. A discrete set of infection-regulated genes responded strongly (>2.5 fold) to DMKG supplementation, including genes involved in the MAPK signaling pathway and in cytokine signaling (Table S5). Finally, several non-coding RNA were differentially regulated during infection, in a DMKG sensitive manner, including several tRNAs. SAM is a co-factor for post-transcriptional modifications required for tRNA stability (Adami and Bottai, 2020). Thus, SAM depletion during infection (Fig. 3A) may take part in the decrease in tRNA levels. Interestingly, a drop in host protein synthesis was previously reported in *Chlamydia* infected cells (Ohmer et al., 2019). The decrease in tRNA levels could contribute to this phenomenon.

Our data converge to link protein hypermethylation in infection to the metabolic pressure exerted by bacterial growth. Does it benefit the host or the bacteria? The question is difficult to answer in the absence of a complex model in which histone methylation levels could be controlled experimentally. *In vitro*, iron supplementation reduced histone hypermethylation without affecting progeny production, indicating that hypermethylation is not required for the generation of infectious bacteria. Furthermore, it is likely that the bacteria have adapted their growth rate to limit possible deleterious consequence of the metabolic pressure they exert on the host. Interestingly, several demethylase inhibitors hindered bacterial proliferation *in vitro*, indicating that increasing histone methylation beyond its physiological range was deleterious for the bacteria. We tested the effect of one broad demethylase inhibitor, JIB-04, on the outcome of infection in an animal model. A reduction of bacterial burden upon exposure to JIB-04 was observed, although below statistical significance with only 6 animals per group. A trend towards reduced tissue damage was also observed, but the effect remained modest. Interpretation of these observations calls for caution, as exposition to JIB-04 might affect non-infected cells implicated in bacterial clearance. Thus, the trend to a reduction in bacteria burden and pathology might not be related to the decrease in bacterial proliferation upon treatment with demethylase inhibitors observed *in vitro*. Still, these data suggest that chemical modifications of the epigenome of the host can impact the outcome of infection.

In conclusion, the effect of metabolites on the epigenetic status of chromatin are documented in metabolic diseases, in cancer and on the influence exerted by the gut microbiota on host physiology. We show here that the metabolic activity of an intracellular bacterium has the potential to compromise metabolic homeostasis of the host and to weigh on the host response to infection through epigenetic alterations.

## MATERIAL AND METHODS

### Cells

HeLa cells (ATCC) and mouse embryonic fibroblasts (MEFs) were grown in Dulbecco’s modified Eagle’s medium with Glutamax (DMEM, Invitrogen), supplemented with 10 % (v/v) heat-inactivated fetal bovine serum (FBS). Primary epithelial cells were isolated from ectocervix biopsies of female patients after approval by the French Ethical Committee ‘CPP Ile de France 1’ on May 9, 2016. Approval and authorization of the National Data Protection authority (‘Commission Nationale de l’Informatique et des Libertés’, CNIL) have been obtained for the research protocol, in compliance with the Helsinki principles. Informed consent was obtained from all participants. Primary cells were cultivated in keratinocyte serum free medium (KSFM, Thermo Fisher Scientific) containing 50 mg/L of bovine pituitary extract (Thermo Fisher Scientific) and 5 μg/L of epidermal growth factor (EGF) human recombinant (Thermo Fisher Scientific) (Tang *et al*., 2021). They were used between passage 3 and 6. All cell cultures were maintained at 37 °C, in 5 % CO2 atmosphere and were routinely tested for mycoplasma by PCR.

### Bacteria

The *C. trachomatis* serovar LGV L2 strain 434 was from ATCC. For IFU measurements and for image analysis, this strain stably transformed with a plasmid expressing the green fluorescent protein (GFP) or mCherry under the control of the *incD* promoter (L2^IncD^GFP and L2^IncD^mCherry) were used, respectively (Agaisse and Derré, 2013). For the experiments with organoids *C. trachomatis* L2 or *C. muridarum* CM006 expressing GFP in p2TK2 or pNigg derived vectors respectively were used (Dolat and Valdivia, 2021). For Fig. S1 *C. muridarum* was the MoPn strain provided by Scot Ouellette (University of Nebraska). The *C. trachomatis* serovar LGV L2 strain 434 with an allele in *nuE* leading to a W71* truncation (C to T mutation at position 129786) was obtained by chemical mutagenesis as previously described (Kokes *et al*., 2015). The resulting rifamycin resistant mutant CTL2M340 was back-crossed with a spectinomycin resistant wild type L2 strain to give rise to the CTL2M340rs59 recombinant strain (referred here as Nue*). Nue* counts 16 single nucleotides polymorphisms (SNPs) compared to the parental strain, which are listed in Table S8. Bacteria were propagated on HeLa cells using standard procedures (Scidmore, 2005).

### Immunofluorescence

Cells (50×10^3 per well) were seeded in a 24 well-plate on a glass coverslip and infected at MOI=1 the following day. In some experiments, the medium was changed 24 hpi for complete medium supplemented with ferric ammonium citrate (MP Biomedicals #158040 100 mM stock), or dimethyl α-ketoglutarate (DMKG, Sigma #349631, the 7.18 M preparation was adjusted to 2.5 M stock with DMSO, aliquoted and frozen), or JIB-04 (Sigma #SML0808 5 mM stock in DMSO), or doxycycline (Sigma #D3072 10 μg/ml stock in culture medium), or ciprofloxacin (Sigma 17850 0.3 mM stock in culture medium) or SD70 (Xcess Biosciences #M60194-2S, 40 mM stock in DMSO) or TACH101 (MedChemExpress #HY-O0297 and #HY-O0298 1:1, 150 μM stock in DMSO). Cells were fixed 48 hpi unless otherwise indicated in 4% paraformaldehyde (PFA) 4% sucrose in phosphate buffer saline (PBS) for 20 min, followed with 10 min in 50 mM NH4Cl in PBS, and then washed in PBS. Coverslips were permeabilized with 0.3% Triton-X100 in PBS for 5 min and washed in PBS. Samples were incubated for 1 hour in primary antibodies diluted in 1mg/ml bovine serum albumin (BSA) 0.01% Tween-20 in PBS, washed 3 times in PBS, and then incubated for 1 hour in secondary antibodies with 0.5 μg/ml 4,6-diamidino-2-phenylindole (DAPI) (Invitrogen). Antibodies used are listed in Table S6. Samples were washed in PBS and then mounted on slides in mowiol. Microscopy was performed on an Axio observer Z1 microscope equipped with an ApoTome module (Zeiss, Germany) and a 20X Apochromat lens. Images were taken with an ORCAflash4.OLT camera (Hamamatsu, Japan) using the ZEN software from Zeiss.

### Image analysis and quantification

Image analysis was performed with Fiji (Schindelin et al.), the different steps described below were automatically performed using a Fiji macro script. Cell nuclei and inclusions appeared in the uv/DAPI channel (DNA stain) while only inclusions appeared in the red/mCherry channel (mCherry expressing bacteria). A median filter of radius 2 was applied to both channels for noise removal. “Triangle” and “Huang” automatic threshold methods were respectively used on mCherry and DAPI channels to create binary masks. The mCherry mask was subtracted from the DAPI mask, resulting in a binary mask corresponding to cells nuclei. The FIJI functions “Erode”, “Dilate”, “Fill Holes” and “Watershed” were successively applied to the resulting mask for nuclei individualization. The FIJI functions “Erode”, “Dilate” 2 times and “Watershed” were successively applied to the inclusion mask to individualize inclusions and dilate them by one pixel to create an overlap with the nuclei next to inclusions. Regions of interest (ROI) corresponding to inclusions were defined on the inclusions mask using “Analyze Particles” function with “Size” parameter set to “30-Inf”. ROI corresponding to nuclei were defined on the nuclei mask using “Analyze Particles” function with “Size” parameter set to “50-500” and “Exclude on edges” option on. For each nuclei ROI, the macro checked each inclusion ROI one by one to see if there was an overlap with the nuclei ROI. This was done by selecting the intersection of both ROI (“AND” function of the ROI Manager) and by checking if it was not empty (selectionType() > -1). Nuclei were considered as from an infected cell if they were overlapping with at least one inclusion. For Fig. 2B, the overlap area was measured, and nuclei were associated with the inclusion they were overlapping the most with. The mean intensity in each channel was measured for each nucleus and inclusion. Finally, for each field, three composite pictures of DAPI and mCherry channels were saved, displaying either inclusions, infected or non-infected nuclei segmentation for postprocessing verification. The mean value of intensity for a given histone mark in non-infected samples was calculated. Nuclei whose intensity was greater than two times this value were counted as positive. Positive nuclei from non-infected cells were manually double checked and corrected as nuclei from infected cells if they had been misclassified by the automated procedure.

### Organoid culture and imaging

Murine endometrial organoids were generated as previously described from C57BL/6J (Jackson Laboratory, strain no. 000664) (Dolat and Valdivia, 2021). Bacteria were diluted in phosphate-buffered saline to a final concentration of 5 x 10^6 IFU, vortexed for 15 seconds, and pipetted into a borosilicated capillary glass needle. Organoids were microinjected using an Eppendorf FemtoJet 4x using a steep vertical angle and only punctured once and refed with media containing 10 ug/mL gentamicin one hour after injection. At the indicated timepoint, organoids were washed with warm PBS, fixed with 3% warm paraformaldehyde for 20 minutes, and resuspended in 0.5 mL 0.25% ammonium chloride. Organoids were centrifuged at 600 x *g* for 5 minutes, resuspended in PBS containing 2% BSA (Fraction V; Equitech-Bio) and 0.1% triton X-100 (Sigma), and incubated on a rocker at 25 °C. The organoids were centrifuged at 600 x *g* for 5 min and incubated with rabbit anti-H3K9me3 (1:50) overnight at 4 °C. Organoids were washed once with 2% BSA, centrifuged at 600 x *g* for 5 min, and incubated with anti-rabbit conjugated to AlexFluor 647 (1:1,000) and Hoechst (2 ug/mL) for 1.5 hours at 25 °C. Organoids were centrifuged at 600 x *g* for 5 minutes, resuspended in Vectashield (Vector Labs; H-1000), pipetted onto a glass slide and overlaid with a coverslip. Organoids (9 organoids for infections with *C. trachomatis* and 7 for *C. muridarum*) were imaged using an inverted microscope (TI2-E; Nikon Instruments) equipped with an ORCA Flash 4.0 V3 sCMOS camera (Hamamatsu), SOLA solid-state white illumination (Lumencor) and acquired using a 20x Plan Apochromatic NA 0.75 air objective with the NIS-Elements Software (Nikon). 3D reconstructions were generated in ImageJ (NIH) and the nuclei were segmented using auto local thresholding and the watershed algorithms. The mean intensity in infected cells and neighboring, uninfected cells was measured, plotted and analyzed in R Studio. Statistical analysis was performed using a Welch’s t-test.

### Proteomic analyses

35 mm dishes were plated with 400.000 HeLa cells and infected or not the following day with *C. trachomatis* L2 at MOI=2. Forty-eight hours later the cells were collected and resuspended in Tris 100mM, 8M urea pH 7.5. Biological triplicates were prepared. Proteins were reduced using 5 mM TCEP for 30 min at room temperature. Alkylation of the reduced disulfide bridges was performed using 10 mM iodoacetamide for 30 min at room temperature in the dark. Proteins were then digested in two steps, first with 1 µg r-LysC Mass Spec Grade (Promega) for 4 h at 30°C and then samples were diluted to 1.8 M urea with 100 mM Tris HCl pH 8.5 and 1 µg Sequencing Grade Modified Trypsin was added for the second digestion overnight at 37°C. Proteolysis was stopped by adding formic acid (FA) at a final concentration of 5%. The resulting peptides were desalted on Stage Tip (Rappsilber et al., 2007) prepared with Empore 3M C18 material (Fisher Scientific). Peptides were eluted using 50 % acetonitrile (ACN), 0.1 % formic acid (FA). Peptides were concentrated to dryness and resuspended in 2 % ACN/0.1 % FA just prior to LC-MS injection.

*LC-MS/MS analysis* was performed on a Q Exactive^TM^ Plus Mass Spectrometer (Thermo Fisher Scientific) coupled with a Proxeon EASY-nLC 1200 (Thermo Fisher Scientific). One µg of peptides was injected onto a home-made 32 cm C18 column (1.9 μm particles, 100 Å pore size, ReproSil-Pur Basic C18, Dr. Maisch GmbH, Ammerbuch-Entringen, Germany). Column equilibration and peptide loading were done at 900 bars in buffer A (0.1 % FA). Peptides were separated with a multi-step gradient from 3 to 6 % buffer B (80% ACN, 0.1% FA) in 5 min, 6 to 31 % buffer B in 130 min, 31 to 62 % buffer B in 30 min at a flow rate of 250 nL/min. Column temperature was set to 60°C. MS data were acquired using Xcalibur software using a data-dependent method. MS scans were acquired at a resolution of 70,000 and MS/MS scans (fixed first mass 100 m/z) at a resolution of 17,500. The AGC target and maximum injection time for the survey scans and the MS/MS scans were set to 3E6, 20ms and 1E6, 60ms respectively. An automatic selection of the 10 most intense precursor ions was activated (Top 10) with a 30 s dynamic exclusion. The isolation window was set to 1.6 m/z and normalized collision energy fixed to 27 for HCD fragmentation. We used an underfill ratio of 1.0 % corresponding to an intensity threshold of 1.7E5. Unassigned precursor ion charge states as well as 1, 7, 8 and >8 charged states were rejected and peptide match was disable.

#### Protein identification

Acquired Raw data were analyzed using MaxQuant software version 1.6.6.0 using the Andromeda search engine (Tyanova et al., 2016). The MS/MS spectra were searched against the *Chlamydia trachomatis* serovar L2-434-Bu and the Homo Sapiens Uniprot reference proteome database (884 and 79057 entries respectively). All searches were performed with oxidation of methionine, mono- and di-methylation of lysine and arginine, trimethylation of lysine, and acetylation of lysine and protein N-terminal as variable modifications and cysteine carbamidomethylation as fixed modification. Trypsin was selected as protease allowing for up to two missed cleavages. The minimum peptide length was set to 5 amino acids and the peptide mass was limited to a maximum of 8,000 Da. One unique peptide to the protein group was required for the protein identification. The main search peptide tolerance was set to 4.5 ppm and to 20 ppm for the MS/MS match tolerance. Second peptides was enabled to identify co-fragmentation events and match between runs option was selected with a match time window of 0.7 min over an alignment time window of 20 min. The false discovery rate (FDR) for peptide and protein identification was set to 0.01. The mass spectrometry proteomics data have been deposited to the ProteomeXchange Consortium via the PRIDE partner repository with the dataset identifier PXD051857.

*Statistical analysis* on proteomics data was based on the comparison of intensities between biological conditions. Proteins identified as reverse hits or potential contaminants were first removed from the analysis. Intensities associated with at least one unique peptide were kept for further statistics. Additionally, only proteins with at least two intensity values in one of both conditions were kept to ensure a minimum of replicability. Next, intensities were log2-transformed. They were normalized so that the intensities of human proteins are centered in each sample on the mean of the medians of the human protein intensities across all samples. This normalization was carried out to ensure that the intensities in each sample corresponded to a same global level of the host proteins. Proteins without any intensity value in one of both conditions have been considered as proteins quantitatively present in a condition and absent in the other. They have, therefore, been set aside and considered as differentially abundant proteins. Next, missing values of the remaining proteins were imputed using the impute.mle function of the R package imp4p (Gianetto et al., 2020). Statistical testing was conducted using limma t-tests thanks to the R package limma (Ritchie et al., 2015). The FDR control was performed using an adaptive Benjamini–Hochberg procedure on the resulting p-values thanks to the function adjust.p of R package cp4p using the robust method to estimate the proportion of true null hypotheses among the set of statistical tests (Giai Gianetto et al., 2016). The proteins associated to an absolute log2(fold-change) superior to 1 and adjusted p-values inferior to a FDR level of 1% have been considered as significantly differentially abundant proteins.

#### Statistical analysis of modified peptides

We searched for mono-, di- or tri-methylations on lysine or arginine residues. Some modifications were identified through mass differences in MS1 spectra, without precise localization on the peptide. Others have been pinpointed on their respective peptides using MS/MS spectra with localization probabilities exceeding 0.75 as determined by MaxQuant. For statistical analyses, raw intensities of all identified modified peptides in the experiments were log2-transformed. Subsequently, they were normalized by median centering within the biological conditions (infected or not infected) to account for potential shifts in peptide median signals between conditions. Missing values were then imputed by condition using the impute.mle function from the R package imp4p. Standard t-tests were employed to examine whether the fold-changes of modified peptides significantly differed from 0 between conditions. Furthermore, ANOVA contrast analyses were conducted to identify modified peptides with fold-changes significantly different from those of the unmodified proteins to which they belong as explained in (Giai Gianetto, 2023).

### Nuclei isolation and measure of DNA methylation

35 mm dishes were seeded with 4×10^5 HeLa cells. Cells were infected or not at MOI=2 with *C. trachomatis* LGV L2. Forty-eight hpi, cells were washed in PBS and detached in PBS-EDTA, centrifuged for 5 min at 400 x*g* and resuspended in 344 ml of hypotonic buffer (10 mM Tris pH 7.65, 1.5 mM MgCl2, 10 mM KCl, 0.2% NP-40, 0.6 mM phenylmethanesulfonyl (PMSF), 1X Protease inhibitor cocktail (Sigma)). Volume was adjusted to about 1 ml by adding PBS, cells were then lysed using a Dounce homogenizer (20 up and down), and then 96 μl of Sucrose buffer (20 mM Tris-HCl pH 7.65, 15 mM KCl, 60 mM NaCl, 0.34 M sucrose, 150 mM spermine (Merck S4264), 50 mM spermidine (Merck S0266), 0.6 mM PMSF, 1X Protease inhibitor cocktail) was added to the samples. Samples were centrifuged for 5 min at 500 x*g*, supernatants were discarded and the pellets frozen. Genomic DNA was extracted with RNase A and digested with the kit Nucleoside digestion mix of NEB (M0649S) according to the manufacturer’s protocol. The reaction was processed for LC-MS/MS analysis. Analysis of total 5-methyl-2ʹ-deoxycytidine (5-mdC) concentrations was performed using a Q exactive mass spectrometer (Thermo Fisher Scientific), equipped with an electrospray ionization source (H-ESI II Probe) coupled with an Ultimate 3000 RS HPLC (Thermo Fisher Scientific). Digested DNA was injected onto a Thermo Fisher Hypersil Gold aQ chromatography column (100 mm x 2.1 mm, 1.9 µm particle size) heated at 30 °C. The flow rate was set at 0.3 mL/min and run with an isocratic eluent of 1% acetonitrile in water with 0.1 % formic acid during 10 min. Parent ions were fragmented in positive ion mode with 10 % normalized collision energy in parallel-reaction monitoring (PRM) mode. MS2 resolution was 17,500 with an AGC target of 2e5, a maximum injection time of 50 ms and an isolation window of 1.0 m/z. The inclusion list contained the following masses: dC (228.1), 5-mdC (242.1). Extracted ion chromatograms of base fragments (±5ppm) were used for detection and quantification (112.0506 Da for dC; 126.0662 Da for 5-mdC). Calibration curves were previously generated using synthetic standards in the ranges of 0.2 to 50 pmoles injected for dC and 0.02 to 10 pmoles for 5mdC. Results are expressed as a % of total dC.

### SAM measurement

For each condition, a 10 cm dish was seeded with 2.5 million HeLa cells. Cells were infected or not at MOI=2 with *C. trachomatis* LGV L2. Forty hpi, nuclei were isolated as described above. The samples were divided in 8 aliquots and centrifuged at 500 x*g* for 5 minutes. The pellet (nuclei) was resuspended in 40 ul of CM buffer from the Mediomics Bridge-It® SAM fluorescence assay (Gentaur) (2 aliquots, flash-frozen) or of urea buffer (30 mM Tris, 150 mM NaCl, 8 M urea, 1 % SDS, pH=8.0) (1 aliquot) to determine protein concentration using a BCA protein assay kit (Pierce 23225). Aliquots were thawed and normalized to equal protein concentrations in CM buffer, diluted 1:4 in CM buffer and incubated for 1 h at 24 °C with vigorous shaking. Samples were centrifuged for 5 min at 16,000 x*g*. The supernatant was diluted 1:5 in Buffer S, mixed and 20 ul of the mix was deposited on one Optiplate (Perkin Elmer #6005270) in duplicates. 80 ul of Reaction Buffer was added to each well. The reaction was incubated for 40 min at room temperature before reading fluorescence resonance energy transfer (FRET) on a FLUOStar Omega with filters (485-12 excitation 540nm-10 emission, 485-12 excitation 660-10 emission). SAM concentrations in test samples were then determined using a SAM standard curve.

### H3K4me3 demethylation activity and succinate measurement

10 cm dishes were seeded with 1.5 million HeLa cells (2 dishes were used for the infected plates, to compensate for a lower nucleus harvest). Cells were infected or not at MOI=2 with *C. trachomatis* LGV L2. Forty hpi, nuclei were isolated as described above and resuspended in 0.3 ml of ice-cold PBS with 1X Protease inhibitor cocktail. Samples were sonicated 3 times for 5 s with cooling on ice between sonication steps and centrifuged for 10 min at 16,000 x*g* at 4 °C. The supernatants were collected and centrifuged a second time for 5 min. The supernatants were immediately used to measure their ability to demethylate H3K4me3 by immune-detection of the reaction product using the Epigentek kit P-3083. Briefly, 5 ul of nuclear extracts was mixed to 45 ul of the manufacturer’s assay buffer JD2 supplemented with 2 uM ascorbate, loaded on a plate coated with H3K4me3 and incubated for 120 min at 37 °C. The demethylated products were revealed with a specific antibody following the manufacturer’s instructions and fluorometrically measured on a plate reader (Tecan) at 530 nm excitation and 590 nm emission. Background readings were given by heat inactivation of the nuclear extracts prior to the demethylase reaction (15 min at 85 °C). The remainder of the nuclear extracts were flash-frozen and later used to determine protein and succinate concentrations using BCA and Succinate-Glo™ JmjC Demethylase/Hydroxylase (Promega V7990) assay kits, respectively. Nuclear extracts were heat inactivated prior to succinate measurement, samples were diluted 1:10 and 25 uL were used. Protein concentrations were used to normalize H3K4me3 demethylase activity and succinate concentrations in nuclear extracts.

### Progeny assays

Hela cells (10^5) were seeded in a 24-well plate. The next day cells were infected with L2^IncD^GFP bacteria at a MOI=0.15. Twenty-for post infection, the medium was replaced with fresh medium supplemented with the indicated drug, or left unchanged, with the addition of DMSO in the well if the drug was dissolved in DMSO. Cells were detached 48 hpi, lysed using glass beads and the supernatant was used to infect new HeLa cells plated the day before (100 000 cells/well in a 24-well plate), in serial dilution. The next day, 3 wells per condition with an infection lower than 30 % (checked by microscopy) were detached and fixed as described above, before analysis by flow cytometry and determination of the bacterial titer. Acquisition was performed using a CytoFLEX S (Beckman Coulter) and 50 000 events per sample were acquired and then analyzed using FlowJo (version 10.0.7).

### Transfections

One day after being seeded in a 24 well-plate (10^5 cells/well) in 500 μL of complete medium transfection mix was added to cells (50 μL of Jetprime buffer (Polyplus), 0.5 μg of DNA, and 3 μL of Jetprime transfection reagent (Polyplus) per well. After 3h, the medium was changed for new complete medium containing mCherry-bacteria at MOI=0.5. Plasmids used were coding for HA-tagged KDM4A/JMJD2A, KDM5A and dKDM5A (H483A mutation) were obtained from Addgene (#24180, #14800 and #14801 respectively). The plasmid coding for dKDM4A was generated from the wild type construct by restriction-free cloning (van den Ent and Löwe, 2006) using primers ctcgacggatcaattcaccat and aggtagttgatgctgtagaggtccatgtcttcagtgGCccaagcaaaggatgtcttcc. This resulted in His to Alanine mutation in the catalytic site (position 188).

### Chromatin immunoprecipitation

Four 15 cm culture dishes were seeded with 6×10^6 cells) and half were infected the day after at MOI=2. Forty-eight h later cells were rinsed twice in PBS and fixed by adding 15 ml of fixation medium prewarmed to 37 °C (15 mM NaCl, 0.15 mM EDTA, 0.075 mM EGTA, 0.015 mM HEPES in DMEM pH8.0. Formaldehyde (Sigma F-8775) was added to the prewarmed medium just prior to the fixation step to the final concentration of 1%. Cells were rocked gently at room temperature, afterwards formaldehyde was quenched by adding glycine drop by drop from a 1.06M stock prepared fresh to a finale concentration of 0.125 M. Quenching was pursued for 5 min with gentle rocking, the cells were rinsed twice with PBS, scraped using a soft silicon scraper and transferred to a 15 ml conical tubes on ice, pooling the two identical plates together. Cells were centrifuged at 400 x*g* for 10 min at 4 °C, the pellet was resuspended in 10 ml cold PBS, centrifuged again. The pellet was resuspended in 1 ml of cold PBS and centrifuged, the supernatant was discarded and the pellet frozen at – 80 °C. Pelets were resuspended in lysis buffer (50 mM HEPES (pH 7.5), 140 mM NaCl, 1 mM EDTA (pH 8), 10% glycerol, 0.5% NP-40, 0.25% Triton X100) and incubated 10 min at 4°C. After centrifugation (300 x g, 5 min, 4°C) the supernatant was discarded, the pellet resuspended in buffer (200 mM NaCl, 1 mM EDTA (pH 8), 0.5 mM EGTA (pH 8), 10 mM Tris-HCl (pH 8) and FPIC (Fast Protease Inhibitors Cocktail; Sigma-Aldrich; #S8830-20TAB) and incubated 10 min at 4 °C. After centrifugation the supernatant was discarded and the pellet resuspended in low-SDS buffer (50 mM Tris-HCl (pH 8), 0.1% SDS, 0.95% NP-40, 0.1% NaDoC, 10 mM EDTA (pH 8), 150 mM NaCl and FPIC) for sonication for 4 min (4 x (30 s ON, 30 s OFF)) (Bioruptor, Diagenode), yielding genomic DNA fragments with a bulk size of 150–500 bp. 1% of chromatin extracts were taken aside for inputs. Chromatin corresponding to 5 μg of DNA was incubated with 2 μg of anti-H3K4me3 (Diagenode; Cat#: C15410003) or anti-H3K9me3 (Diagenode; Cat#: C15410193) antibodies overnight at 4◦C. The DNA-protein-antibody complexes were captured by DiaMag protein A-coated magnetic beads (Diagenode; Cat#: C03010020) by incubating at 4°C for 2 h. Magnetic beads were then washed with low-salt buffer (20 mM Tris-HCl (pH 8), 0.1% SDS, 1% Triton X100, 2 mM EDTA (pH 8), 150 mM NaCI and FPIC) and high-salt buffer (20 mM Tris–HCl (pH 8), 0.1% SDS, 1% Triton X100, 2 mM EDTA (pH 8), 500 mM NaCI and FPIC). DNA was eluted with TE buffer (100 mM Tris-HCl (pH 8), 1% SDS, 1 mM EDTA (pH 8)) from the beads overnight, and then reverse cross-linked with 1 μl of RNase A (10 μg/ml; Sigma-Aldrich; Cat#: R4642), followed by Proteinase K digestion (5 μl of 20 mg/mL; Sigma-Aldrich; Cat#: P2308) during 2 h; inputs corresponding to 1% of IP were reverse cross-linked in the same conditions. DNA was purified using MinElute PCR Purification Kit (Qiagen; Cat#: 28004). 2 μl of input and ChIP were loaded on High Sensitivity D5000 ScreenTape (Agilent Technologies #5067-5592) with 2 μl of reagent (#5067-5593) (Fig. S4A). Libraries were prepared using the MicroPlex Library Preparation Kit v3 (Diagenode) according to the manufacturer’s protocol. In brief, 1 to 25 ng were used as input material. After adapters ligation, fragments were amplified with Illumina primers for 10 cycles. Libraries were purified using AMPure XP beads protocol (Beckman Coulter, Indianapolis, IN) and quantified using the Qubit fluorometer (Thermo Fisher Scientific). Libraries’ size distribution was assessed using the Bioanalyzer High Sensitivity DNA chip (Agilent Technologies). Libraries were normalized and pooled at 2 nM and spike-in PhiX Control v3 (Illumina) was added. Sequencing was done using a NextSeq 500 instrument (Illumina) in paired-end mode (2×40 cycles) and a NextSeq 2000 instrument (Illumina) in paired-end mode (2×50 cycles). After sequencing, a primary analysis based on AOZAN software (ENS, Paris, France) (Perrin et al., 2017) was applied to demultiplex and control the quality of the raw data (based on bcl2fastq v2.20, bcl-convert v4.1.5 and FastQC modules / v0.11.5).

ChIP-seq datasets were analyzed under the Galaxy platform (https://usegalaxy.eu) (The Galaxy, 2022). The Quality of raw data has been evaluated with FastQC (Galaxy Version 0.74) (https://www.bioinformatics.babraham.ac.uk/projects/fastqc/). Poor quality sequences and adapters have been trimmed or removed with Trimmomatic (Galaxy Version 0.23.2) software (Bolger et al., 2014) to retain only paired reads of 40 bp with Phred score >20. Reads were mapped to the reference genomes (GRCh38) using Bowtie2 (Galaxy Version 2.5.0) (Langmead et al., 2019) to generate BAM files. Duplicates were removed by using the Picard tool MarkDuplicates (Galaxy Version 2.18.2.4). Narrow and broad peaks were called with MACS2 (Galaxy Version 2.2.7.1) (Feng et al., 2012) using the input as control, FDR ≤ 0.05. Differential binding analysis of ChIP-Seq peak data was performed using DiffBind (Galaxy Version 2.10.0) (Ross-Innes et al., 2012). Peak annotation was performed by using the ChIPseeker R package (Yu et al., 2015) and Bedtools intersect (Galaxy Version 2.28.0) (Quinlan and Hall, 2010) to annotate peaks according to their chromatin feature with genome segmentation data from ENCODE (Dunham et al., 2012; Ernst and Kellis, 2012). The Gene ontology enrichment analysis was performed with the clusterProfiler R package (Yu et al., 2012). The significant GO categories were identified with Benjamini-Hochberg adjusted p-value ≤ 0.05. Data can be found under the GEO accession number GSE265792.

### Microarray

A 6-well plate was seeded with 200,000 HeLa cells per well. The following day, cells were infected with *C. trachomatis* LGV L2 at MOI=2. 24 hpi, the media was replaced by complete medium supplemented with 5 mM DMKG or left unchanged with addition of an equal volume of DMSO. Forty-eight hpi RNA was harvested using the RNeasy kit (Qiagen), and their quality was analyzed using the BioAnalyzer (Agilent). Transcriptome analysis was done using the GeneChip Human Transcriptome Array 2.0 for in combination with the GeneChip WT PLUS Reagent kit (Applied Biosystems). 33,3ng total RNA was used as input for the analysis that was done according to the manufacturer’s guidelines. In brief, double-stranded cDNA is first synthesized using random primers and then used as template for in vitro transcription (IVT) that produces amplified amounts of antisense mRNA (cRNA). The cRNA is used as input for a second round of first strand cDNA synthesis, producing single stranded sense cDNA. After fragmentation and end-labeling the targets are hybridized to single sample GeneChip cartridge arrays that are stained and washed on a GeneChip Fluidics Station 450 and scanned on a GeneChip scanner 3000 7G. Cell files containing the probe cell intensity data generated by the scanner is imported into the Expression Console Software 4.0.1 (Thermo Fisher Scientific) for data analysis. Expression values corresponds to Robust Multiarray Average (RMA) normalized probe set intensities. Data can be found under the GEO accession number GSE265791.

### RT-qPCR and qPCR

Total RNAs from cells in culture were isolated 48 hpi with the RNeasy Mini Kit (Qiagen) with DNase treatment (DNase I, Roche). RNA concentrations were determined with a spectrophotometer NanoDrop (Thermo Fisher Scientific) and normalized to equal contents. Reverse transcription (RT) was performed using the M-MLV Reverse Transcriptase (Promega). Quantitative PCR (qPCR) was undertaken on DNA or on complementary DNA (cDNA) with LightCycler 480 system using LightCycler 480 SYBR Green Master I (Roche). Data were analyzed using the ΔΔCt method with the *actin* gene as a control gene (Schmittgen and Livak, 2008). Each RT-qPCR experiment was performed in duplicate and repeated at least three times. Primer used are listed in Table S7.

### tRNA quantification

tRNA were extracted and analyzed by northern blot as described (Torres et al., 2019) with the exception that in some cases samples were run on a 6% PAGE (instead of 8%) to get better separation between the 5S rRNA and the U6 snRNA.

### Infection in mice

The experiments were performed in accordance with the French national and European laws regarding the protection of animals used for experimental and other scientific purposes. The study was approved by the ethics committee of Institut Pasteur (approval N° dap210086). Female c57BL/6J mice aged 6-7 weeks were purchased from Charles River Laboratories (France) and housed in the animal facility at Institut Pasteur. JIB-04 was synthesized as described (Wang *et al*., 2013). All animals were treated with 2.5 mg of medroxyprogesterone (Depo-provera-SC®, Pfizer) 7 days prior to infection to synchronize their menstrual cycle. Vaginal lavages were conducted the day prior to infection twice with 50 μl PBS each time by pipetting up and down several times followed with swabbing to remove the mucus. On day 0, the animals were anesthetized by intramuscular injection of Ketamine/Xylazine in PBS before intra-peritoneal (i.p.) injection of JIB-04 (100 mg/kg body weight) resuspended in 300 μl sesame oil (Sigma #3547), control animals were injected with sesame oil only. 2×10^5^ IFUs of *C. muridarum* resuspended in 5 μl of sucrose-phosphate-glutamic acid buffer (SPG: 10 mM sodium phosphate [8 mM Na2HPO4-2 mM NaH2PO4], 220 mM sucrose, 0.50 mM l-glutamic acid) were introduced in the vagina of mice. Additional doses of JIB04 or sesame oil were injected i.p. at D2 and D4. A vaginal lavage was performed at D4 and two washes were pooled and frozen immediately into liquid nitrogen. Total RNAs and genomic DNA from lavage fluids were extracted using AllPrep® DNA/RNA Micro Kit (Qiagen, #80284). Upon D35 the animals were sacrificed by cervical dislocation. The genital tracts were excised, imaged and their morphology examined. Each image was segmented using Icy (https://icy.bioimageanalysis.org/) (de Chaumont et al., 2012) based on color quantization. From the binary image, a distance map and skeleton of the uterine horns were computed using the plugin Morphomath. The distance maps, corresponding to the radius of each uterus horn, were computed starting from the point of convergence of the two uterine horn skeletons.

### Statistical analyses

The R environment v4.1.2 was used for analysis of tRNA quantification and RT-qPCR (R Core Team, 2019). Data were not averaged prior analysis. For tRNA quantification, band intensities were measured in each well of northern blot pictures and the log2 of (band of interest intensity / control band intensity) was computed, with band of interest being either GlyGCC or ArgGCU tRNA, and with control band being either U6 or 5S rRNA. Then, data were fitted to a mixed linear model that included the condition and the technical replicates as random covariate (i.e., band intensity ∼ condition + (1|replicate)). For RT-qPCR analysis, a variation of the ΔΔCt method (gene of interest versus actin) (Schmittgen and Livak, 2008) that keeps the technical replicate variations was applied. These values were fitted to a mixed linear model that included the treatment (DMSO, DMKG), the infection (Infected, Not infected) and the technical replicates as random covariate (i.e., ΔΔCt2 ∼ treatment * infection + (1|replicate) in Fig 5C). Contrasts were performed with the emmeans() function of the emmeans package. Statistical significance was set to P ≤ 0.05. In each panel, type I error was controlled by correcting the p values according to the Benjamini & Hochberg method (“BH” option in the p.adjust() function of R). The nature of the statistical tests used in other figures are provided in the respective legend.

## ACKNOWLEDGEMENTS

We thank Béatrice Niragire, Klementina Borovnik and Valentin Petit for technical assistance, Drs MS Phan and JY Tinevez for the pipeline of analysis of images of infected cells with non-fluorescent inclusions, Drs Mélanie Hamon, Paola Arimondo, Sophie Polo and Mikhail D. Magnitov for discussion and reagents. CIC was supported by the Pasteur - Paris University (PPU) International PhD Program. This work was supported by the Agence Nationale pour la Recherche 20-PAMR-0011 TheraEPI, the Fondation ARC pour la recherche sur le Cancer, the Institut national du cancer INCA_16719, the Institut Pasteur, the Centre National de la Recherche Scientifique, the Welch foundation (grant I-1878 to EDM).

---

**Fig. S1.**
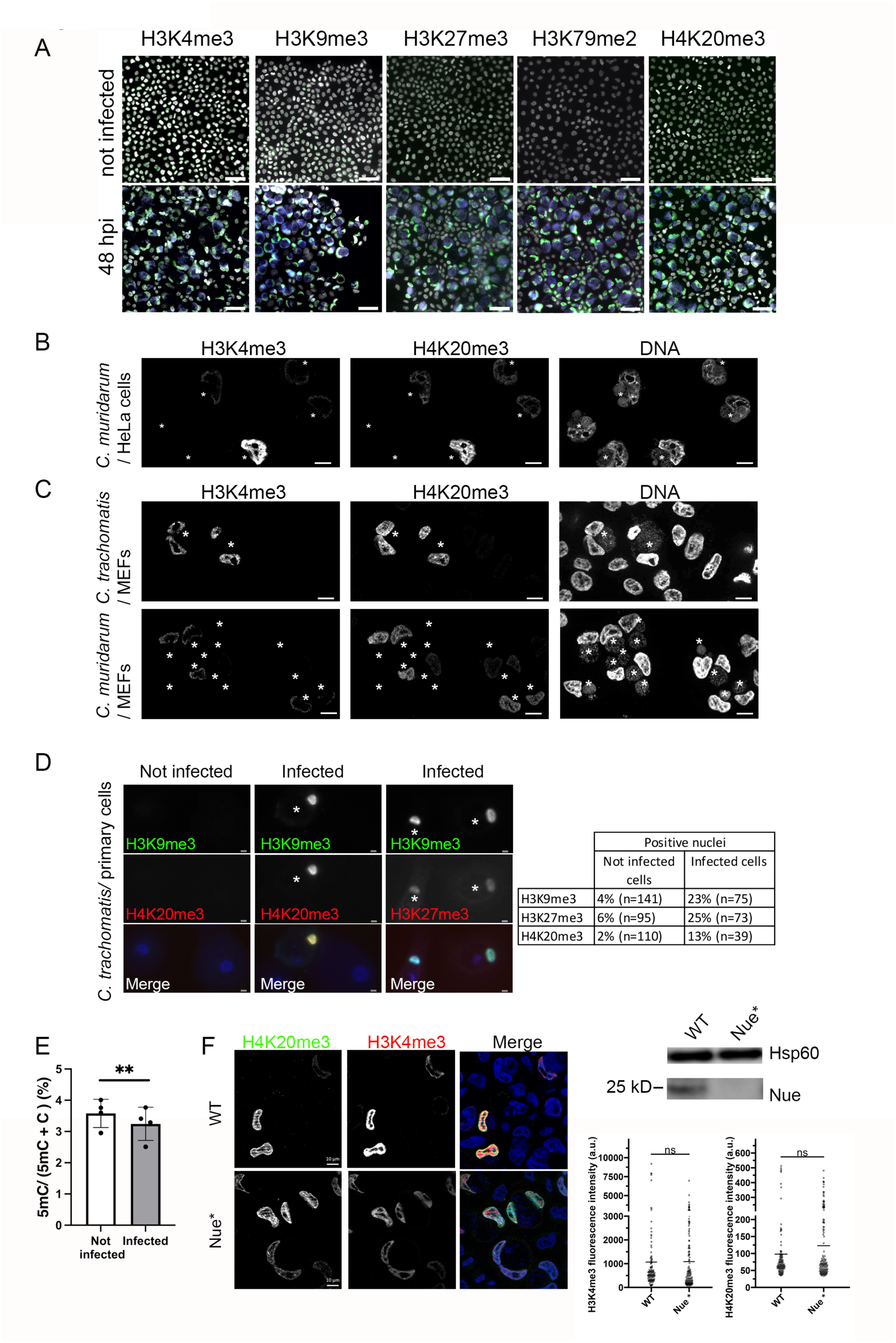
Histone hypermethylation occurs in various cell type and is independent of Nue. **A**. HeLa cells were infected or not with mCherry expressing *C. trachomatis* L2 for 48 h before fixation, permeabilization and staining with antibodies specific for the indicated histone methylation marks, followed with Alexa-488 conjugated secondary antibodies (green). DNA was labeled with DAPI (white), mCherry is displayed in blue. Bar, 50 μm. **B**. HeLa cells were infected with *C. muridarum* for 48 h before fixation, permeabilization and staining for H3K4me3 and H4K20me3. DNA was stained with DAPI, inclusions are highlighted with an asterisk. **C**. MEFs were infected with *C. trachomatis* L2 (top) or *C. muridarum* (bottom) for 48 h before proceeding as in B. **D**. Primary human epithelial cells isolated from the ectocervix were infected for 48 h with *C. trachomatis* serovar L2 before proceeding as in B. Right panel shows quantification of relative distribution of histone marks. **E**. Nuclei were isolated from cells infected for 48 h with *C. trachomatis* L2 or from control cells prior to measurement of DNA methylation by mass spectrometry. Results of four independent experiments are shown, with mean ± SD, and the P-value of the Student’s paired t-test is indicated (**P < 0.01). **F**. Cells were infected for 48 h with *C. trachomatis* L2 WT or with a mutant that does not express Nue (Nue*=CTL2M340rs59, see methods), fixed, permeabilized and stained with the indicated antibodies. The dot plots display the intensity for the indicated histone methylation marks in the nuclei of infected cells. Absence of Nue expression in cell lysates 24 hpi was verified by western blot. Antibodies against the chlamydial protein Hsp60 were used to control for infection levels.

**Fig. S2.**
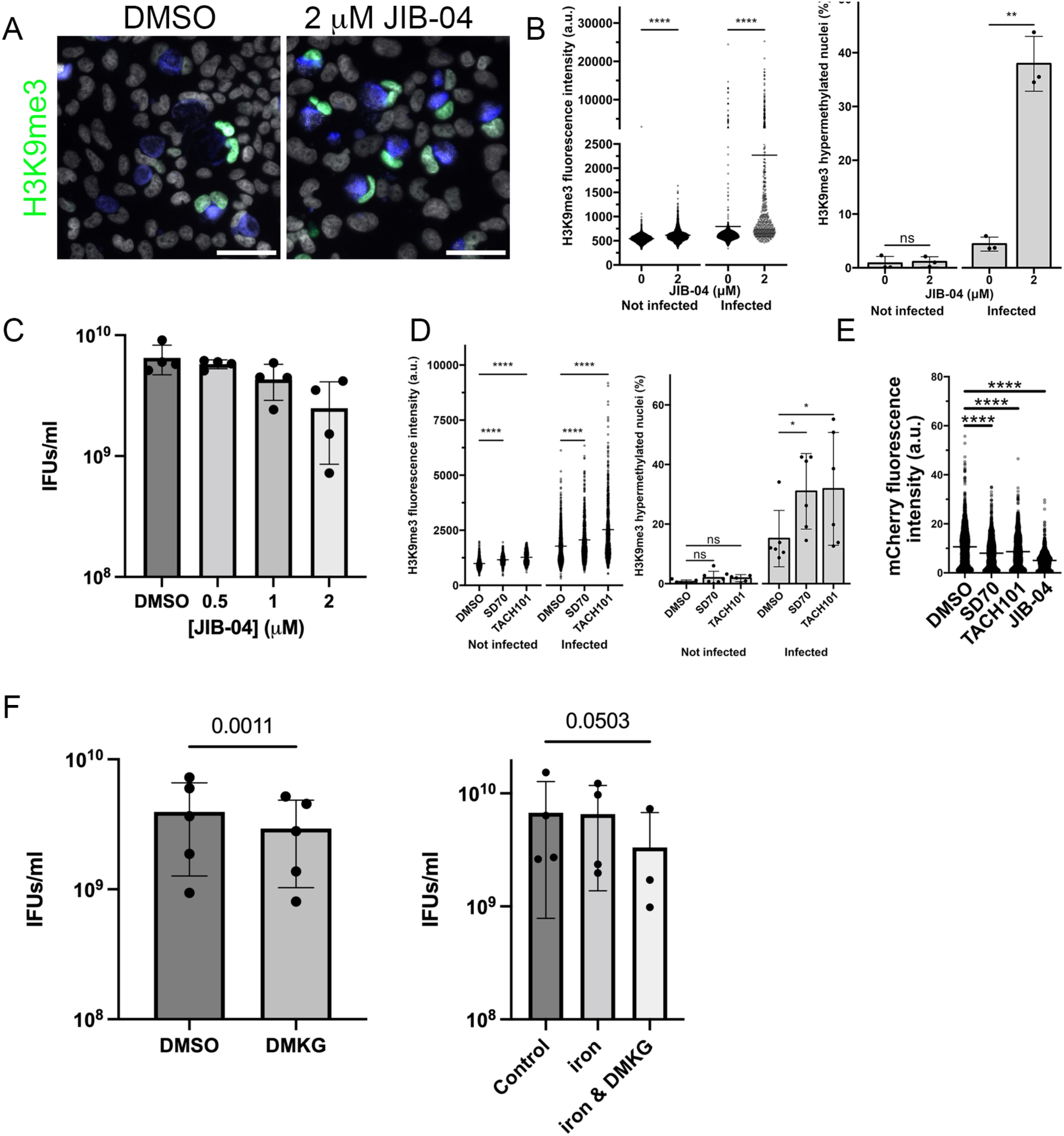
Effect of KDM inhibitors, DMKG or iron on histone methylation and on infection. **A**. Cells were infected with mCherry expressing *C. trachomatis* serovar L2. Twenty-for hours later the culture medium was renewed with or without 2 μM JIB-04. Cells were fixed 48 hpi, permeabilized and stained for H3K9me3 (green). DNA was labeled with DAPI (white), mCherry is displayed in blue. Bar, 50 μm. **B** H3K9me3 mean fluorescence intensity in nuclei of non-infected or infected cells treated 24 hpi with the indicated concentration of JIB-04 and fixed and stained 48 hpi in one representative experiment. a.u., arbitrary unit. Black bars indicate the mean. Histograms display the percentage of positive nuclei in three independent experiments. The p-values of Welch’s t-tests are shown ****, P < 0.0001; **, P < 0.01. **C**. HeLa cells were infected with *C. trachomatis* L2 expressing GFP at MOI=0.2 and treated with the indicated concentration of JIB-04 24 hpi. Infectious forming units (IFUs) collected 48 hpi were quantified by reinfecting fresh HeLa cells as described in the methods. **D**. The KDM4 inhibitors SD70 (2.5 μM), TACH101 (10 nM) or vehicle (DMSO 1:1000) were added in the culture medium 24 hpi. Samples were processed 48 hpi and analyzed as in B. **E**. The mCherry intensity in the inclusions in the images analyzed in D is used as an indicator of bacterial load, the effect of JIB-04 (2 μM) is shown for comparison. **F**. Same as C, with replacement of the culture medium with fresh medium complemented with 5 mM DMKG (left), 100 μM ferric ammonium citrate or a combination of the two (right). The result of three to five independent experiments and the p-values of ratio-paired t-tests are shown.

**Fig. S3.**
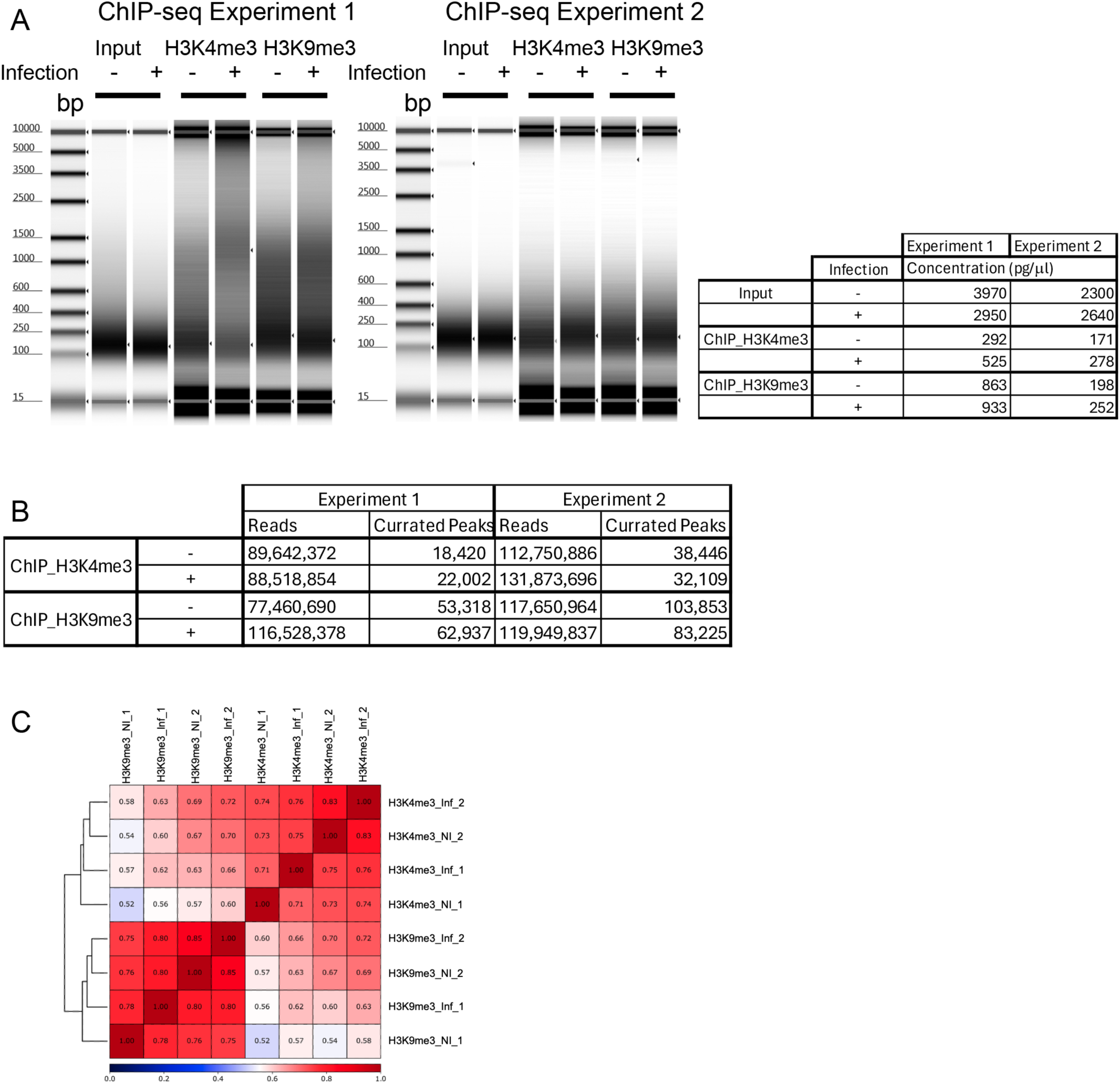
Quality controls for the ChIP experiments. **A**. Electophoresis profile of input DNA isolated from infected and uninfected cells and corresponding ChIP with antibodies to the indicated histone marks. The DNA concentrations measured in each lane are reported in the table. **B**. Number of reads and peaks analyzed for each experiment. **C**. Hierarchical clustering of correlation matrix of H3K4me3 and H3K9me3 ChIP-seq replicates (1 and 2) performed in non-infected (Nl) and infected (Infected) HeLa cells. Spearman correlations were calculated in deepTools (the multiBamSummary (Galaxy Version 3.5.4+galaxy0) was followed with plotCorrelation (Galaxy Version 3.5.4+galaxy0) using the read counts split into 500-bp bins across the human genome.

**Table S1.** Relative abundance of host methylated proteins 48 hpi with *C. trachomatis*.

**Table S2.** Relative abundance of host proteins 48 hpi with *C. trachomatis*.

**Table S3.** ChIP-seq analyses.

**Table S4.** Differentially expressed host genes 48 hpi with *C. trachomatis*.

**Table S5.** DMKG sensitive genes in *C. trachomatis* infection.

**Table S6.** Antibodies used.

|  | Target | Species | Supplier | Catalog # |
| --- | --- | --- | --- | --- |
| Primary Antibodies | H3K9me3 | Rabbit | Abcam | ab8898 |
|  | H4K20me3 | Mouse | Abcam | ab78517 |
|  | H3K27me3 | Mouse | Bp biosciences | 25243 |
|  | H3K4me3 | Rabbit | Diagenode | C15410003 |
|  | H3K79me2 | Rabbit | Abcam | ab3594 |
|  | HA | Rat | Roche Diagnostics | Clone 3F10 #1 867 423 |
|  | Nue | Rabbit | Subtil lab | Pennni et al 2010 |
|  | Hsp60 | Rabbit | Subtil lab |  |
| Secondary Antibodies | Alexa Fluor TM 488 goat anti-rabbit IgG (H+L) | Goat | Invitrogen | A-11034 |
|  | Alexa Fluor TM 647 goat anti-rabbit IgG (H+L) | Goat | Invitrogen | A-21244 |
|  | Alexa Fluor TM 488 goat anti-mouse IgG (H+L) | Goat | Invitrogen | A-11029 |
|  | Alexa Fluor TM 647 goat anti-mouse IgG (H+L) | Goat | Invitrogen | A-21235 |
|  | Alexa Fluor TM 488 goat anti-rat IgG (H+L) | Goat | Invitrogen | A-11006 |

**Table S7.**
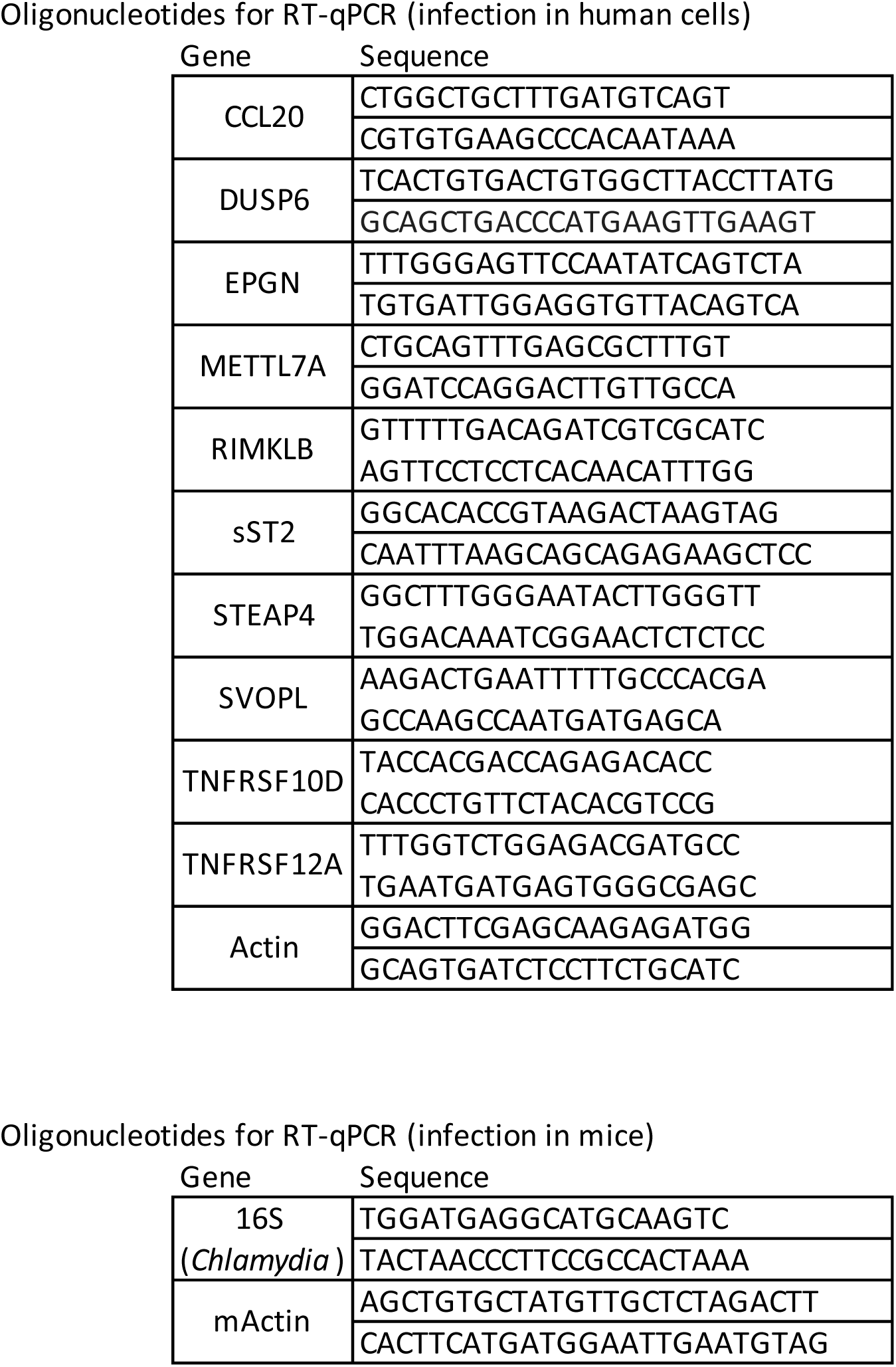
Oligonucleotides used.

**Table S8.**
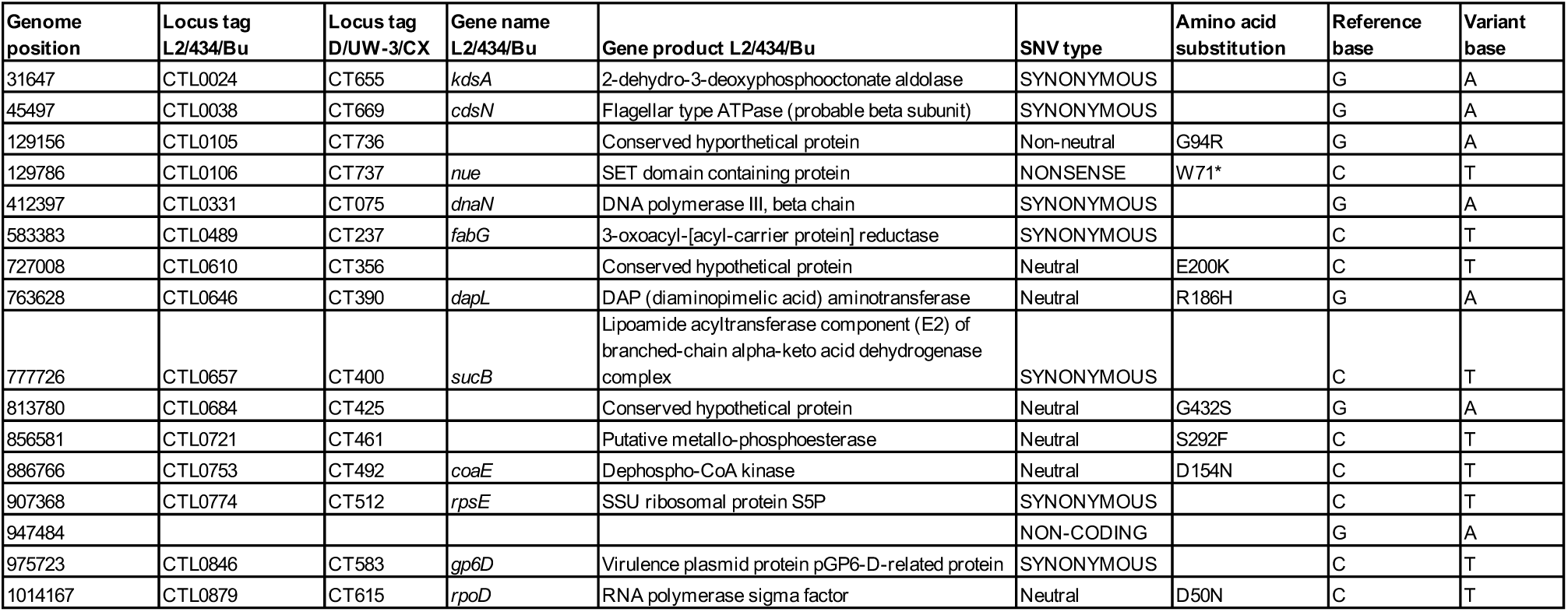
CTL2M340rs59 variants, identified by genotyping.

| Genome position | Locus tag L2/434/Bu | Locus tag D/UW-3/CX | Gene name L2/434/Bu | Gene product L2/434/Bu | SNV type | Amino acid substitution | Reference base | Variant base |
| --- | --- | --- | --- | --- | --- | --- | --- | --- |
| 31647 | CTL0024 | CT655 | *kdsA* | 2-dehydro-3-deoxyphosphooctonate aldolase | SYNONYMOUS |  | G | A |
| 45497 | CTL0038 | CT669 | *cdsN* | Flagellar type ATPase (probable beta subunit) | SYNONYMOUS |  | G | A |
| 129156 | CTL0105 | CT736 |  | Conserved hypothetical protein | Non-neutral | G94R | G | A |
| 129786 | CTL0106 | CT737 | *nue* | SET domain containing protein | NONSENSE | W71* | C | T |
| 412397 | CTL0331 | CT075 | *dnaN* | DNA polymerase III, beta chain | SYNONYMOUS |  | G | A |
| 583383 | CTL0489 | CT237 | *fabG* | 3-oxoacyl-[acyl-carrier protein] reductase | SYNONYMOUS |  | C | T |
| 727008 | CTL0610 | CT356 |  | Conserved hypothetical protein | Neutral | E200K | C | T |
| 763628 | CTL0646 | CT390 | *dapL* | DAP (diaminopimelic acid) aminotransferase | Neutral | R186H | G | A |
| 777726 | CTL0657 | CT400 | *sucB* | Lipoamide acyltransferase component (E2) of branched-chain alpha-keto acid dehydrogenase complex | SYNONYMOUS |  | C | T |
| 813780 | CTL0684 | CT425 |  | Conserved hypothetical protein | Neutral | G432S | G | A |
| 856581 | CTL0721 | CT461 |  | Putative metallo-phosphoesterase | Neutral | S292F | C | T |
| 886766 | CTL0753 | CT492 | *coaE* | Dephospho-CoA kinase | Neutral | D154N | C | T |
| 907368 | CTL0774 | CT512 | *rpsE* | SSU ribosomal protein S5P | SYNONYMOUS |  | C | T |
| 947484 |  |  |  |  | NON-CODING |  | G | A |
| 975723 | CTL0846 | CT583 | *gp6D* | Virulence plasmid protein pGP6-D-related protein | SYNONYMOUS |  | C | T |
| 1014167 | CTL0879 | CT615 | *rpoD* | RNA polymerase sigma factor | Neutral | D50N | C | T |

